# Capturing R-loops at base-pair resolution unveils clustered R-loops regulate gene expression in a number-dependent manner

**DOI:** 10.1101/2025.02.21.639597

**Authors:** Yaoyi Li, Yingliang Sheng, Chao Di, Hongjie Yao

## Abstract

R-loops are prevalent triplex nucleic strands found across various organisms, involved in numerous biological processes. However, the physiological and pathological functions of R-loops remain largely unknown due to a lack of effective and high-resolution detection methods. Here, using nuclease P1, T5 exonuclease, and lambda exonuclease mediated digestion of ssRNA, ssDNA, and dsDNA while preserving RNA:DNA hybrid, we report a method named R-loop identification assisted by nuclease and high-throughput sequencing (RIAN-seq) for genome-wide mapping of R-loops at base-pair resolution. RIAN-seq represents ultra-accuracy in position and size of R-loops and identifies an order of magnitude more R-loops than current methods. Notably, we find the majority of R-loops ranging from 60 bp to 130 bp and reveal previously unresolvable patterns of R-loops aggerated in clusters across the genome. Clustered R-loops at gene promoters recruit zinc finger transcription factors (VEZF1 and SP5) to promote transcription. The number of R-loops within a cluster positively correlates with the diversity of bound transcription factors. Furthermore, clustered R-loops are less susceptible to transcription perturbation as the number of R-loops within clusters increases. Overall, we demonstrate the ability to identify R-loops at unprecedented resolution and facilitate investigating the mechanisms of clustered R-loops in gene regulation in diverse biological processes.

## Introduction

R-loops are three-stranded nucleic acid structures composed of a DNA:RNA hybrid and the associated non-template single-stranded DNA, existing throughout various species(Sanz et al., 2016; Wahba et al., 2016; Xu et al., 2017) and extensively involved in physiological and pathological processes(García-Muse and Aguilera, 2019). As a regulator of epigenetic functioning(Costantino and Koshland, 2015; Sanz et al., 2016), R-loops have essential functions in regulating gene expression(Skourti-Stathaki et al., 2019), chromatin organization(Luo et al., 2022; Wulfridge et al., 2023; Zhang et al., 2023), and cell fate decisions(Li et al., 2020), underscoring the importance of accurate R-loop detection and characterization. Several methods have been developed to improve our understanding of R-loops at the molecular level, but each has limitations. DRIP-seq, which relies on the S9.6 antibody, was the first developed and most widely used for identifying R-loops(Ginno et al., 2012). DRIP-seq variants, including RDIP(Nadel et al., 2015), DRIPc-seq(Sanz et al., 2016), S1-DRIP-seq(Wahba et al., 2016), ssDRIP-seq(Xu et al., 2017), and bisDRIP-seq(Dumelie and Jaffrey, 2017), have been subsequently developed. However, S9.6-based methods suffer from limited resolution and bias towards GC-rich sequences(Bou-Nader et al., 2022; Chen et al., 2022). Alternative methods, such as R-ChIP(Chen et al., 2017), MapR(Yan et al., 2019), and R-loop CUT&Tag(Wang et al., 2021), have been developed to overcome the bias caused by S9.6-based methods using the hybrid binding domain (HBD) of RNase H, which resolves the DNA-RNA hybrid(Gaidamakov et al., 2005). However, HBD-based methods are less efficient in R-loop identification and often exhibit other biases, such as preferential detection of R-loops in promoters(Chen et al., 2022). Additionally, the detection of R-loops can be compromised by the promiscuous binding of both the S9.6 antibody and HBD of RNase H. The antibody-free method spKAS-seq labels ssDNA in R-loops using N3-kethoxal(Wu et al., 2022). However, the labeling of N3-kethoxal might be hindered by proteins bound to ssDNA in R-loops. Other noncanonical DNA structures containing ssDNA strands(Bansal et al., 2022), such as D-loops, cruciform structures, and DNA hairpins, also hinder accurate R-loop detection by spKAS-seq. The current reported methods have complicated workflows and are intensive with respect to hands-on time. Notably, the inconsistency of R-loop annotation among these methods, in which most of the R-loops captured in one method are missing in other methods, hinders the study of R-loop biology. Therefore, developing a method for precise and sensitive mapping of genome-wide R-loops is desirable.

In this study, we developed an antibody-free and nuclease-assisted high-throughput sequencing technology (RIAN-seq) for R-loops identification at base-pair resolution. RIAN-seq preserves the persistence of RNA:DNA hybrids in R-loops by digestion of ssRNA, ssDNA, and dsDNA with nuclease P1, T5 exonuclease, and Lambda exonuclease. We identified approximately 0.5 million R-loops in HEK293T cells using RIAN-seq, an order of magnitude more than existing methods. Most importantly, we determined that approximately 30% of R-loops form clusters, which are characterized by higher signal, longer size, and higher GC base constitutions. The genes with clustered R-loops exhibit significant downregulation in expression after resolving R-loops via RNase H1, compared to limited changes in the expression of genes with non-clustered R-loops. The more R-loops within clusters at gene promoters were resolved, the more significant downregulation of target genes was. Moreover, clustered R-loops at the promoter act as regulatory elements and recruit transcription factors, such as VEZF1 and SP5, for gene transcription. Our study suggests that RIAN-seq is an unbiased method for identifying R-loops at base-pair resolution and uncovering the unknown functions of R-loops within clusters.

## Results

### Working principles and development of RIAN-seq

To develop a strategy to digest single-stranded nucleic acids (ssRNA, ssDNA) and double-stranded DNA (dsDNA) while maintaining RNA:DNA hybrids, we selected a combination of nucleases: Nuclease P1 to digest ssRNA and ssDNA, T5 exonuclease to resolve ssDNA and dsDNA, and Lambda exonuclease to degrade supercoiled dsDNA and enable conversion of dsDNA to ssDNA (Figure 1A). Specifically, the RIAN-seq protocol involved four main steps: (1) Extraction of genomic DNA from cells; (2) DNA fragmentation with restriction enzymes (Alu I, Dde I, Mbo I, Mse I, the restriction sites of which were widely distributed in mammalian genome), and subsequently treated with Nuclease P1, T5 exonuclease, and Lambda exonuclease; (3) Integration of adaptors to RNA:DNA hybrids using Tn5 transposase; (4) Library construction with a simple one-step PCR and sequencing (Fig. 1a). RIAN-seq has several potential advantages: (1) The simplicity of streamlined protocol minimizes potential sample loss and enables identification of more R-loops. (2) The utilization of Tn5 transposase in library construction enhances data quality and reduces sequencing costs. (3) RIAN-seq does not exhibit bias towards R-loops in specific genomic regions by eliminating affinity enrichment or chemical labels. (4) Nuclease P1, T5 exonuclease, and Lambda exonuclease allow for precise determination of position and size for R-loops, achieving base-pair resolution. By integrating these features, RIAN-seq aims to provide a more efficient and accurate approach for genome-wide R-loop detection.

**Figure 1.**
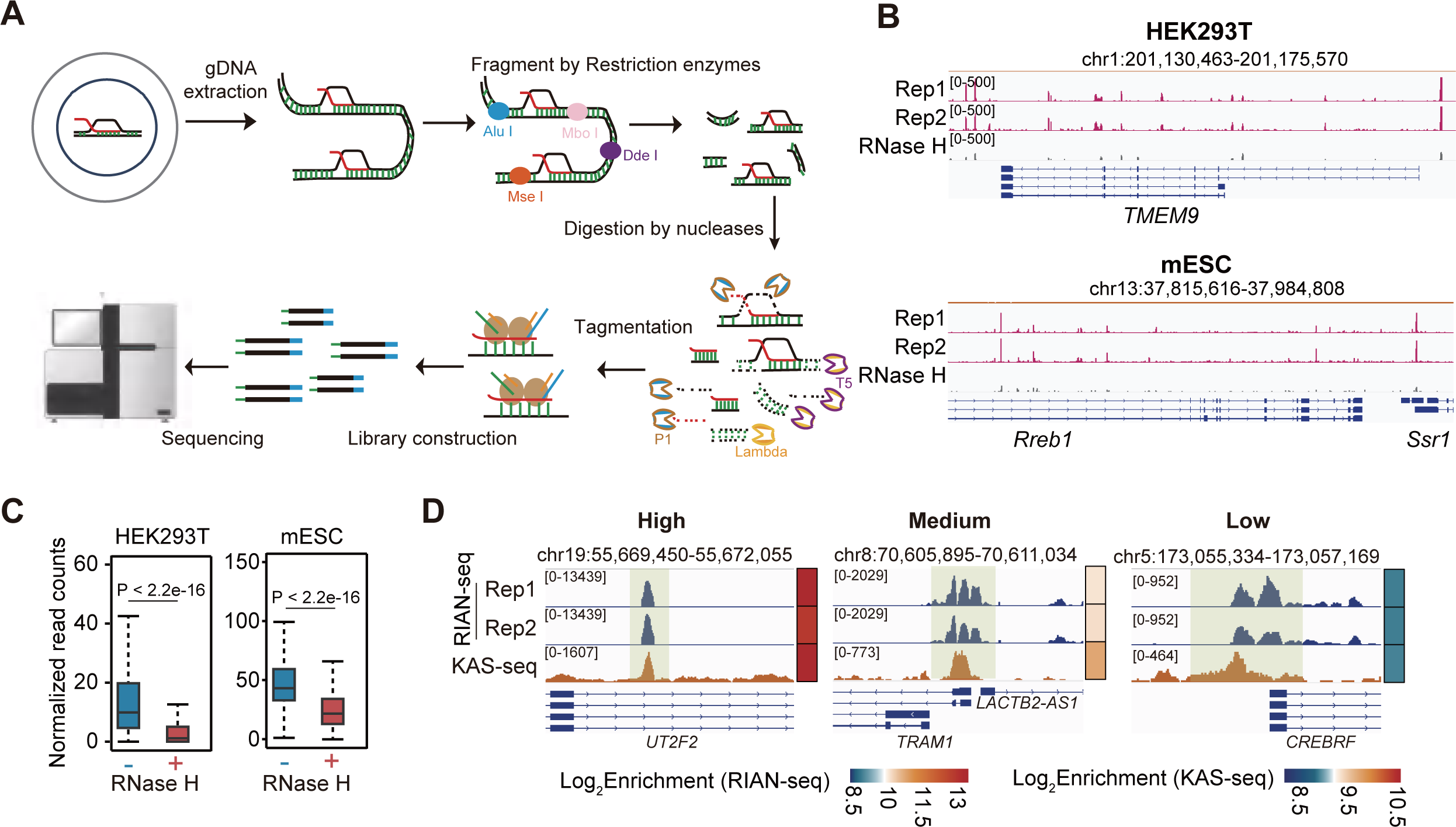
The work strategy and development of RIAN-seq. **A.** The strategic approach of RIAN-seq. P1, Nuclease P1. T5, T5 exonuclease. Lambda, Lambda exonuclease. **B.** Genome browser snapshots depicting the signal of RIAN-seq with and without treatment of RNase H in HEK293T and mES cells, normalized by spike-in. **C.** The box plots showing the signal density of R-loops identified using RIAN-seq treated with or without RNase H in HEK293T cell, normalized by spike-in. **D.** Snapshots indicating high, medium, and low RIAN-seq signals alongside KAS-seq signals in HEK293T cells.

### RIAN-seq faithfully identifies R-loops at the genome-wide scale

To assess the reliability and reproducibility of RIAN-seq, we generated RIAN-seq libraries from HEK293T and mES cells. Two replicates of RIAN-seq in both cell types were highly correlated (Figure S1A and S1B), indicating the reliability of this method. RIAN-seq exhibited a stronger enrichment of signals surrounding transcription start sites (TSSs) (Figure S1C), aligned with the reported distribution of R-loop signals(Petermann et al., 2022). Moreover, approximately 80% of R-loops were shared between biological replicates (Figure S1D), indicating that RIAN-seq enables the generation of highly reproducible datasets between independent biological replicates.

To verify the authenticity of RIAN-seq signals for R-loop detection, we employed treatment with different doses of RNase H during the nuclease digestion step of the RIAN-seq protocol. Our results showed a dosage-dependent reduction in RIAN-seq signals upon RNase H treatment (Figure S2A and S2B). This inverse relationship between RNase H concentration and signal strength indicates that RIAN-seq accurately captures R-loops. To enable quantitative comparisons of samples with or without RNase H-treatment, we added *Drosophila* genomic DNA as a spike-in for internal normalization. As anticipated, RNase H treatment decreased both the overall signals and the number of peaks (Figure 1B, 1C, S2C and S2D), further supporting the specificity of RIAN-seq for R-loop detection. Additionally, RIAN-seq peaks were enriched with signals of KAS-seq(Wu et al., 2020), a method for mapping ssDNA, which is a characteristic structure of R-loops (Figure 1D,S2E and S2F). The high correlation between RIAN-seq and KAS-seq further indicates the accuracy of RIAN-seq in identifying R-loops. Collectively, these findings demonstrate that RIAN-seq reliably and specifically detects R-loops.

### Features and distributions of R-loops detected by RIAN-seq

We then characterized the genomic distribution of R-loops identified by RIAN-seq and observed that R-loops were evenly enriched at promoters (22.04% in HEK293T and 30.68% in mES cells), introns (41.58% in HEK293T and 31.95% in mES cells), and intergenic regions (30.99% in HEK293T and 29.19% in mES cells) (Figure S3A and S3B), all of which represent hotspots for R-loop formation(Chen et al., 2022; Jenjaroenpun et al., 2017; Lin et al., 2022), suggesting that RIAN-seq detects R-loops across various chromatin regions without apparent bias. Moreover, R-loops were strongly correlated with GC and AT skews (Figure S3C and S3D), areas where R-loops frequently occur.

Next, we assessed the relationship between the density of R-loops and its distributions. Firstly, we divided R-loops into three categories based on signal density. The signal of R-loops was highly positively correlated with the level of ssDNA (Figure 1D and Figure S3E), indicating the high fidelity of RIAN-seq in identifying R-loops. Further analysis revealed that the number of R-loops in each signal category was relatively evenly distributed (Figure S3F). These findings demonstrate that RIAN-seq provides a comprehensive and unbiased view of R-loop distribution, capturing R-loops with different signal intensities across the genome.

### RIAN-seq achieves base-pair resolution of R-loops

We examined R-loop distribution in RIAN-seq relative to PRO-seq signals representing active transcription(Kwak et al., 2013). RIAN-seq signals showed strong and sharp enrichment within 500 bp of PRO-seq peaks, contrasting to low or broad distributions flanking 1 kb to 2 kb of PRO-seq peaks in other R-loop detection methods (Figure S4A). Higher RIAN-seq enrichment correlated with regions with higher PRO-seq signals (Figure S4B). RIAN-seq signal strength decreased with increasing distance from PRO-seq peaks (Figure S4C and S4D).

As nucleases could degrade dsDNA located at both sides of R-loops to reduce fragment lengths of the RIAN-seq library, we supposed that RIAN-seq potentially improves the resolution of R-loops. As anticipated, our data indicated that fragment sizes of the RIAN-seq library were predominantly smaller than 100 bp, in contrast to >200 bp fragments in previous methods (Figure S4E). Compared with most R-loops that were >1000 bp in previous methods(Chen et al., 2022), R-loops identified by RIAN-seq primarily ranged from 60 bp to 130 bp (Figure S4F), indicating that RIAN-seq achieves base-pair precision in identifying R-loops (Figure S4G). Furthermore, the size range of R-loops in RIAN-seq is also aligned with direct observations by native electron microscopy and single-molecule imaging studies(Kim et al., 2024; Stoy et al., 2023). These data collectively demonstrate that RIAN-seq substantially improves the resolution of R-loops and provides a more detailed and accurate location and size of R-loops.

### Absolute quantification of R-loops using RIAN-seq

To further evaluate the specificity and sensitivity of RIAN-seq, we conducted a series of controlled experiments using synthetic nucleic acids: synthetic dsDNA to mimic background, synthetic RNA:DNA hybrid to mimic R-loop signals, synthetic ssDNA (the same sequence as dsDNA) and ssRNA (complementary to ssDNA) to assess whether experimental conditions of RIAN-seq gave rise to artifact formation of RNA:DNA hybrid. Synthetic dsDNA was mixed with synthetic RNA:DNA hybrid at various ratios to obtain different RNA:DNA hybrid levels (100%, 60%, 40%, and 0%), accompanied by synthetic ssDNA and ssRNA, among which different nucleic acids were assessed by quantitative PCR (qPCR (Figure S5A)). Our data showed that only synthetic RNA:DNA hybrid was detected (Figure S5A). We observed high agreement (R = 0.998) between expected and experimentally observed RNA:DNA hybrid levels (Figure S5B). Together, these data support the high sensitivity and accuracy of RIAN-seq in R-loop detection.

Additionally, we assessed the signal-to-noise ratios of RIAN-seq at two ratios of RNA:DNA hybrid to dsDNA (3:2, 2:3). qPCR analysis revealed > 0.4 million-fold enrichment of RNA:DNA hybrid signals relative to dsDNA background (Figure S5C), indicating an exceptional signal-to-noise ratio of RIAN-seq. Furthermore, RIAN-seq caused little changes to RNA:DNA hybrid levels (Figure S5D), while the dsDNA background was less than 1/10,000 in contrast to RNA:DNA hybrid (Figure S5A and S5C). These results collectively demonstrate that RIAN-seq offers high specificity and accurate quantification of R-loop levels, achieving an ultra-high signal-to-noise ratio.

### RIAN-seq generates high-quality R-loop profiles at a low sequencing depth

We next investigated the performance of RIAN-seq at different sequencing depths to assess its potential for cost-effective R-loop profiling. RIAN-seq with only 5 million (M) sequencing reads exhibited strong signal enrichment and high correlation (correlation coefficient > 0.9) with results from 100 M reads (Figure S6A and S6B), suggesting that RIAN-seq can provide reliable R-loop profiles even at low sequencing depths. R-loop profiles using just 5 M reads could have similar data quality compared to higher (up to 100 M) (Figure S6B and S6C). Moreover, RIAN-seq showed much sharper signal enrichment at low sequencing depths compared with no clearly visible signal enrichment in previous methods under similar conditions (Figure S6C). Notably, R-loop identification was not saturated at the current maximum sequencing depth (Figure S6D and S6E), and the number of R-loops was still underestimated, indicating the potential for discovering additional R-loops with increased sequencing depth. Thus, we conclude that RIAN-seq is a robust and efficient method for genome-wide R-loop identification.

### RIAN-seq captures an order of magnitude more R-loops than previous methods

We next compared RIAN-seq with published data from other methods, including S9.6-based methods such as DRIP-seq(Ginno et al., 2012) and DRIPc-seq(Sanz et al., 2016), HBD-based methods, such as R-ChIP(Chen et al., 2017), MapR(Yan et al., 2019) and R-loop CUT&Tag(Wang et al., 2021), and chemical label method spKAS-seq(Wu et al., 2022). All methods showed a more robust enrichment of R-loop signals around TSSs compared to other chromatin regions (Figure S7A). R-loops with different signal densities and previously unannotated R-loops in previous methods were identified by RIAN-seq (Figure 2A and S7B). R-loops in RIAN-seq showed individual and sharp signals rather than broad R-loops in DRIP-seq (Figure 2A). Moreover, RIAN-seq showed overall higher signal enrichment (Figure S7C) and more concentrated distribution around ssDNA regions (Figure S7D). These data suggest that RIAN-seq outperforms previous methods in the sensitive and precise profile of R-loops.

**Figure 2.**
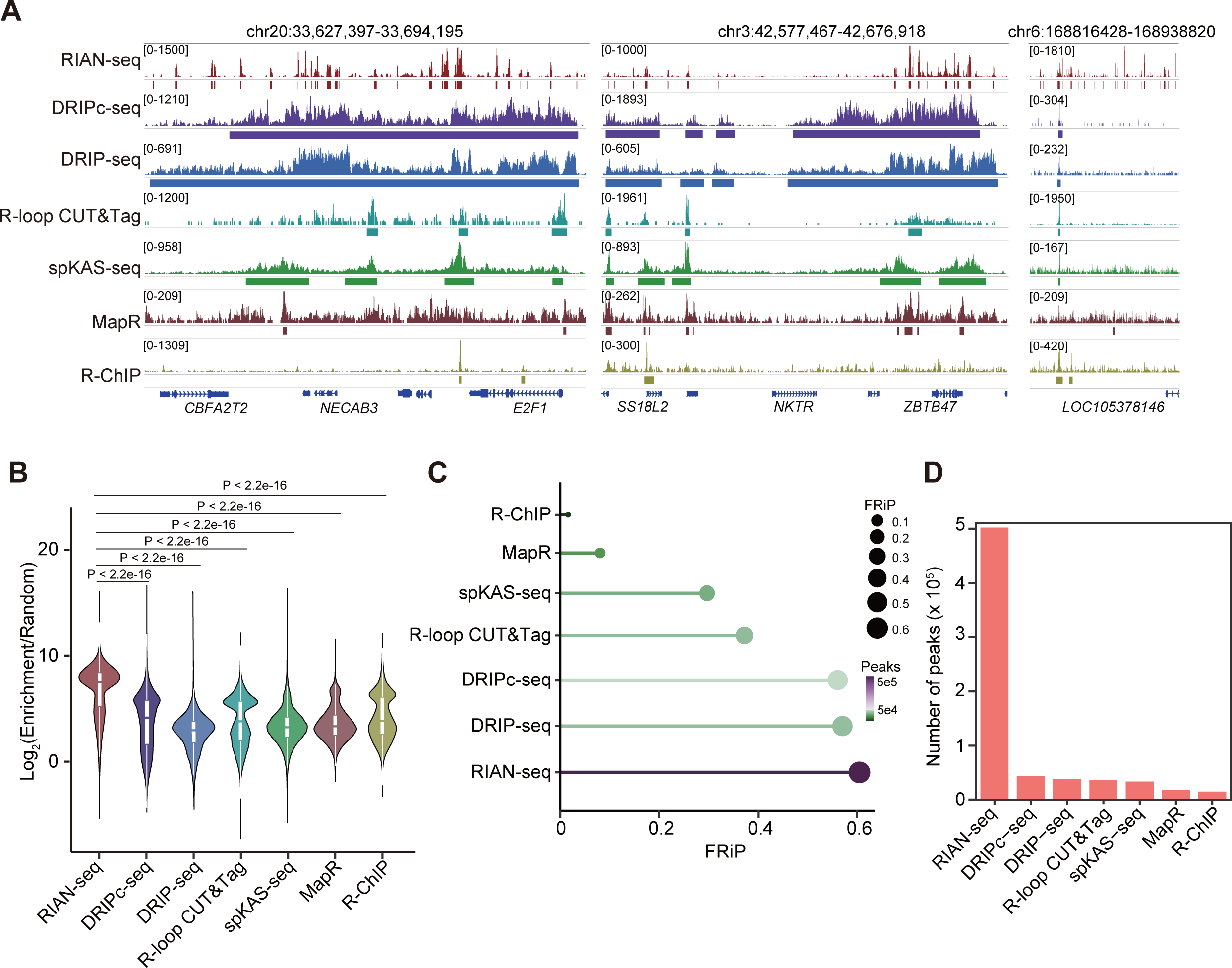
RIAN-seq captures an order magnitude more R-loops than current methods. **A.** Genome browser snapshots depicting the signal and peak distribution of RIAN-seq and previous methods. **B.** A comparison of the signal enrichment of RIAN-seq and previous methods. **C.** Plot illustrating the fraction of reads in peaks (FRiP) scores of RIAN-seq and previous methods. **D.** Histogram showing the numbers of R-loops identified by various R-loop profiling methods.

Quantitatively, RIAN-seq exhibited the strongest enriched signals of R-loops (Figure 2B), with comparable or higher signal-to-noise ratio to previous methods (Figure S7E and S7F). Consistent with the lower background, the most reads falling in peaks of RIAN-seq were determined by the fraction of reads in peaks (FRiP) (Figure 2C). Unexpectedly, RIAN-seq identified approximately 0.5 million R-loops in HEK293T cells (Figure 2D), an order of magnitude more than previous methods (0.01 to 0.08 million R-loops)(Chen et al., 2022).

Due to different enrichment strategies and variable width of R-loops among previous methods, the distribution of R-loops between previous methods is inconsistent, and the majority of R-loops in one method could not be captured by another method(Chen et al., 2022). Excitingly, RIAN-seq captured the majority of R-loops identified by each previous method (Figure S8A). Overlapped R-loops between RIAN-seq and previous methods tended to have higher ssDNA signals (Figure S8B). Besides, RIAN-seq uniquely identified R-loops with lower ssDNA signals, which could not be captured by previous methods (Figure S8B), indicating that RIAN-seq enables the identification of R-loops with low signals.

After combing the R-loops from all other methods, we observed that RIAN-seq captured approximately 60% of combined R-loops identified in all selected previous methods, representing only 40% of RIAN-seq’s total R-loops (Figure S8C). The shared R-loops exhibited higher signals than non-overlapped R-loops (Figure S8D). Consistently, the enrichment of ssDNA signals of overlapped R-loops was much higher than non-overlapped R-loops (Figure S8E). We found that about 80% of overlapped R-loops between RIAN-seq and others were located at promoters and introns, since previous methods are biased in identifying R-loops with high signal at the promoter and gene body(Chen et al., 2022). Besides, our data showed that RIAN-seq unique R-loops were enriched in introns and intergenic regions, suggesting that RIAN-seq has no region bias (Figure S8F). Unique RIAN-seq R-loops at introns and intergenic regions exhibited enrichment in repeat elements, especially SINEs (Figure S8G). Ultimately, RIAN-seq demonstrates superior sensitivity and resolution in R-loop detection, capturing an order of magnitude more R-loops than previous methods, providing a more comprehensive view of R-loop distribution in the genome.

### R-loops form clusters instead of broad peaks

Compared to 1 to 3 R-loops within a gene observed in previous methods, we interestingly found that RIAN-seq identified approximately 10 R-loops within a gene (Figure 2A and S9A), with a fraction of genes possessing over 50 R-loops (Figure 3A). Unlike broad R-loops identified by DRIP-seq distributed throughout the gene, R-loops identified by RIAN-seq were relative narrow. Instead, they showed a tendency to aggregate in clusters (Figure 2A), which is in line with the observation in native electron microscopy analysis(Kim et al., 2024).

**Figure 3.**
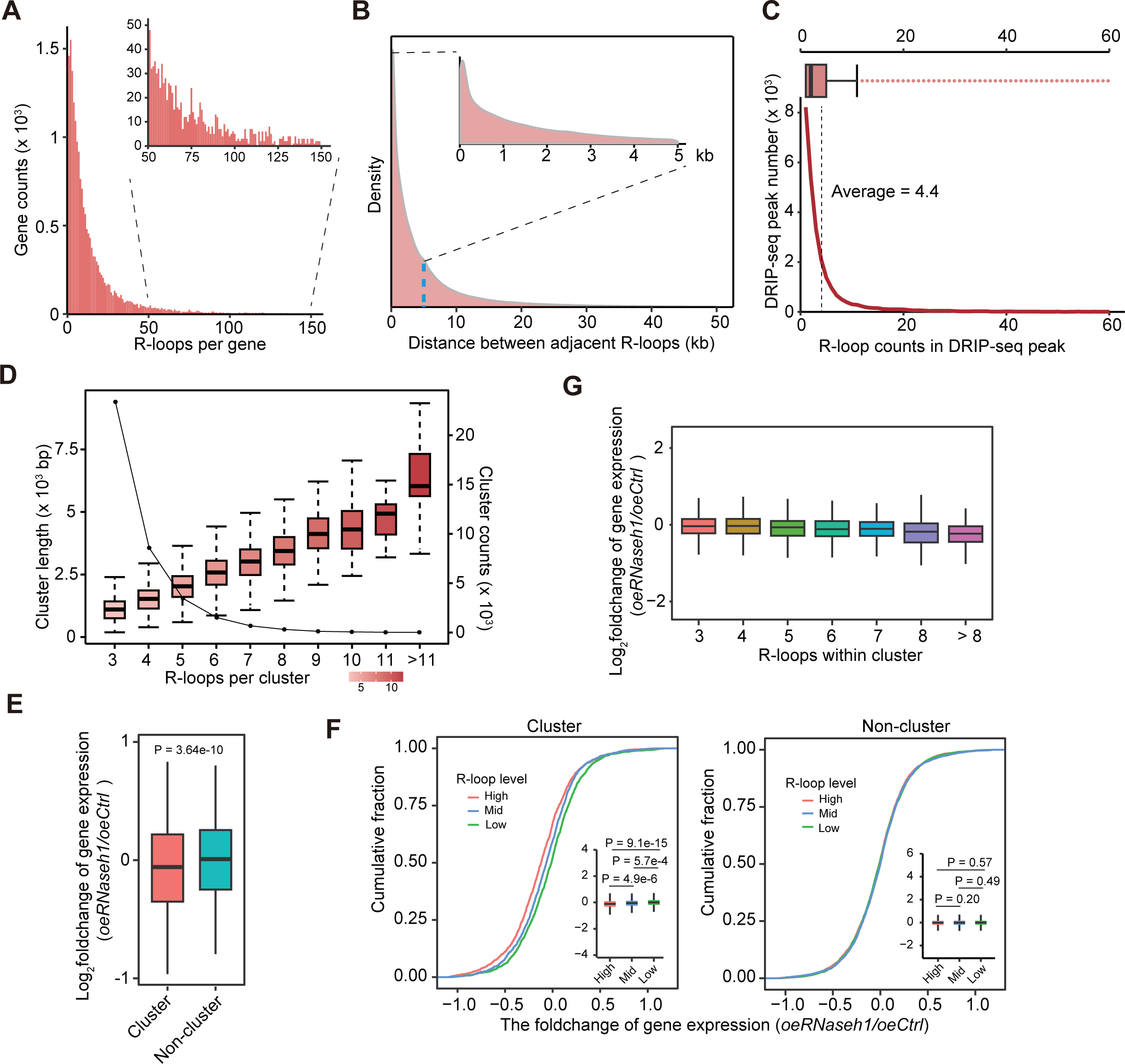
RIAN-seq identifies clustered R-loops not visible in previous methods. **A.** Frequency of genes with R-loop counts from RIAN-seq. **B.** The distributions of two adjacent R-loops within different distances in HEK293T cells. **C.** Curve showing that R-loop peaks detected by DRIP-seq possess one or more RIAN-seq-detected R-loops. Boxes represent the 25th to 75th percentile. **D.** Curve and box plot showing the frequency and length of the clusters containing different R-loop counts. Boxes represent the 25th to 75th percentile. **E.** The boxplots showing the foldchange in the expression of genes with clustered R-loops and non-clustered R-loops after RNaseh1 overexpression. **F.** Cumulative distribution plot of foldchange in genes with clustered and non-clustered R-loops according to R-loop levels. **G.** The boxplots showing foldchange on expression of genes with different numbers of R-loops within a cluster.

To assess whether such clustered R-loops were widespread throughout the genome, we found that adjacent R-loops were significantly enriched within 2 kb regions (Figure 3B), with an average of 4.4 R-loops identified by RIAN-seq within a single DRIP-seq peak (Figure 3C). Focusing on genome-wide R-loop clusters, we observed that three or more adjacent R-loops prone to localize within regions of 3 kb, with the highest proportion of clustered R-loops occurring in approximately 1.5 kb regions (Figure 3D). Using the definition of more than three R-loops within 1.5 kb as an R-loop cluster, we identified 38,167 R-loop clusters including 140,398 R-loops, accounting for 27.6% of R-loops in the genome (Figure S9B). The identified R-loop clusters covered approximately 0.2-9.3 kb, containing 3-22 individual R-loops (Figure 3D). Together, these findings highlight the distinct distribution of R-loops, with a substantial proportion organized into clusters in the genome.

### Clustered R-loops with high GC content are longer than non-clustered R-loops

Next, we explored whether clustered R-loops exhibit distinct characteristics compared to non-clustered R-loops. Clustered R-loops were distributed at promoter, intron, and intergenic regions, while non-clustered R-loops showed a preference for intron and intergenic regions but less at promoters (Figure S9C), indicating potential functional roles of clustered R-loops at promoters. Clustered R-loops showed a preference for GC-rich sequences (Figure S9D). Since the signal of GC-rich R-loops is higher than non-GC-rich R-loops(Crossley et al., 2020), we observed that the overall signals of clustered R-loops were significantly higher than that of non-clustered R-loops (Figure S9E). We further noted that the signal densities were positively correlated with the number of R-loops within clusters (Figure S9F). Moreover, clustered R-loops have relatively longer length than non-clustered R-loops (Figure S9G). We further found that the genes with clustered R-loops at the promoter exhibit dramatically higher expression than those with non-clustered R-loops at promoters (Figure S9H). These data suggest that clustered R-loops possess distinct characteristics from non-clustered R-loops, implying potential roles in transcriptional regulation.

### Clustered R-loops promote transcription

To investigate the roles of clustered R-loops in transcriptional regulation, we overexpressed *RNaseh1*, which specifically resolves RNA:DNA hybrid(Gaidamakov et al., 2005), to reduce R-loop signals in HEK293T cells. RIAN-seq analysis revealed significantly decreased signals of clustered and non-clustered R-loops after *RNaseh1* overexpression (Figure S10A). Low GC content non-clustered R-loops showed a greater signal reduction than higher GC content clustered R-loops (Figure S9D and S10A). Upon loss of R-loops, genes with clustered R-loops showed downregulated expression (Figure 3E). Importantly, the higher signals of clustered R-loops in a given gene, the higher degrees of reduction in its expression, which was not observed in non-clustered R-loops target genes (Figure 3F). In addition, compared with other genomic regions, the loss of clustered R-loops at promoters resulted in the most significant downregulation in gene expression than that of non-clustered R-loops (Figure S10B). These data collectively indicate that clustered R-loops, particularly those at promoter regions, play a crucial role in promoting transcription.

### The numbers of R-loops within a cluster at promoters are essential for gene activation

We next investigated the impact of R-loop number within clusters on gene expression and observed a positive correlation between the number of R-loops within clusters at promoters and the magnitude of gene expression reduction following *RNaseh1* overexpression (Figure 3G). The genes with all R-loops completely resolved within a cluster showed the most dramatic downregulation, particularly in clusters containing five or more R-loops (Figure 4A).

**Figure 4.**
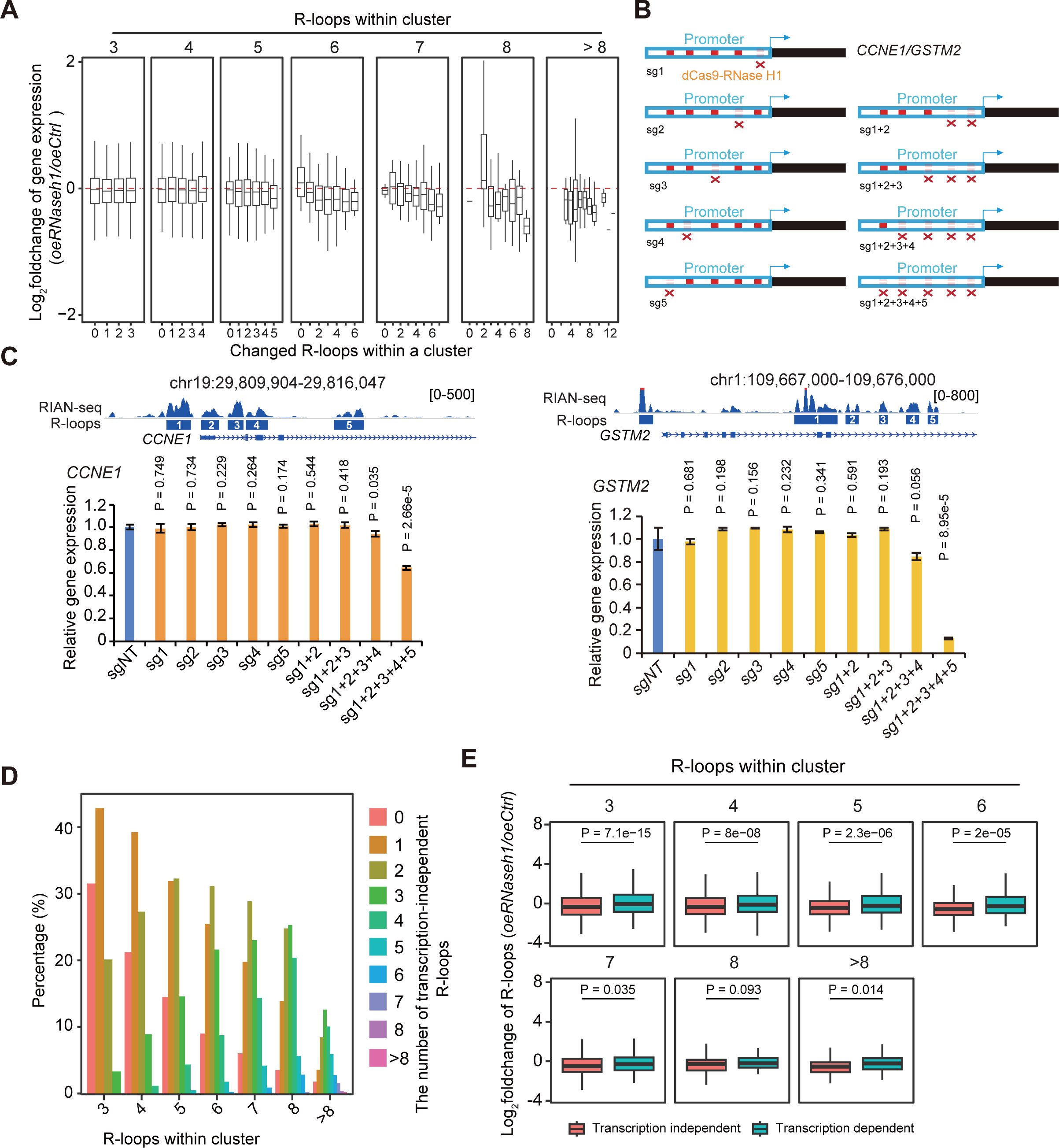
The number of R-loops within a cluster is the primary determinant of transcriptional regulation and the susceptibility of clustered R-loops to transcription perturbation. **A.** Box plots showing the changes in gene expression with different numbers of changed R-loops within a cluster. **B.** The model showing the strategy of site-specific R-loop editing of 5 R-loops clustered at the promoters of both *CCNE1* and *GSTM2*. **C.** The bar graphs showing the expressional changes of *CCNE1* and *GSTM2* with five clustered R-loops at promoters after delivering dCas9-RNase H1 with different combinations of gRNAs targeting R-loops. **D.** The bar graphs showing the frequency of transcription-independent R-loops grouped by the number of R-loops within a cluster. **E.** The comparisons of R-loop signal changes of transcription-dependent and independent R-loop clusters at promoters after *RNaseh1* overexpression.

To investigate whether clustered R-loops coordinately promote transcription, we used a CRISPR dCas9 system fused with RNase H1 (dCas9-RNase H1) for site-specific R-loop resolution (Figure S10C). We delivered single, two, or more guide RNAs targeting clustered R-loops at promoters of the genes *JUP*, *PHACTR4*, *RETREG1*, *CCNE1*, and *GSTM2*, which had 3-5 clustered R-loops at promoters (Figure 4B, S9D and S9E). dCas9-RNase H1 targeting five clustered R-loops at promoters led to greater decreases in expression for both *CCNE1* and *GSTM2*, compared to dCas9-RNase H1 targeting 3 or 4 clustered R-loops at promoters for *JUP*, *PHACTR4,* and *RETREG1* (Figure 4C, S11A and S11B). Moreover, the most significant expression changes of *CCNE1* and *GSTM2* occurred when all clustered R-loops were resolved by dCas9-RNase H1 (Figure 4C and S11B). Therefore, clustered R-loops activate transcription dependent on the number of R-loops within clusters, highlighting the importance of R-loop quantity in transcriptional regulation.

### More R-loops within a cluster are less susceptible to transcription perturbation

To distinguish whether R-loops are transcription-dependent and -independent, we further performed RIAN-seq in HEK293T cells with or without treatments of RNA polymerase II inhibitor, DRB, and Triptolide. Our analysis revealed that approximately 30% of clustered and non-clustered R-loops were unsusceptible to transcription perturbations (Figure S12A). R-loop clusters showed more stability and independent transcription with increased R-loop numbers within a cluster (Figure 4D and S12B). The genes with more transcription-independent R-loops clustered in promoters displayed more significant downregulation of expression after *RNaseh1* overexpression (Figure S12C). Compared to transcription-dependent clustered R-loops, transcription-independent clustered R-loops showed more sensitivity to RNase H1 activity (Figure 4E). Moreover, our data indicated that the R-loop clusters were more prone to be resolved and showed more significant changes in R-loop signals after *RNaseh1* overexpression if the more transcription-independent R-loops existed within a cluster (Figure S12D and S12E). These data indicate that the number of R-loops within a cluster is a key factor in determining the susceptibility of clustered R-loops to transcription perturbation, with more R-loops within clusters being more resistant to such disturbances.

### Clustered R-loops enrich binding motifs of zinc finger transcription factors

To investigate whether any consensus sequences might be present in clustered R-loops, motif analysis indicated that top enriched motifs, RRGRRGVAGG (R = A/G, V =A/C/G), were shown in clustered R-loops, which was different from the motifs of non-clustered R-loops (Figure S13A). Furthermore, approximately 30% of clustered R-loops had the consensus sequence, significantly higher than non-clustered R-loops (Figure S13B). R-loops regulate the chromatin state by recruiting or repelling epigenetic factors(Arab et al., 2019; Chen et al., 2015). Many DNA and RNA binding factors are R-loop-associated proteins(Cristini et al., 2018; Wang et al., 2018), but whether there are any factors for targeting clustered R-loops at promoter regions remains unknown. We aligned the consensus sequence of clustered R-loops with binding motifs of all transcription factors in the database(Vorontsov et al., 2024) and found four zinc finger proteins (VEZF1, SP5, ZNF467, and ZNF341) as candidate factors with high degree of confidence (Figure 5A). More than 50% of clustered R-loops at promoters contained the motifs of these factors (VEZF1, SP5, ZNF467, and ZNF341), with more R-loops within clusters enriching more motifs of these transcription factors (Figure 5B).

**Figure 5.**
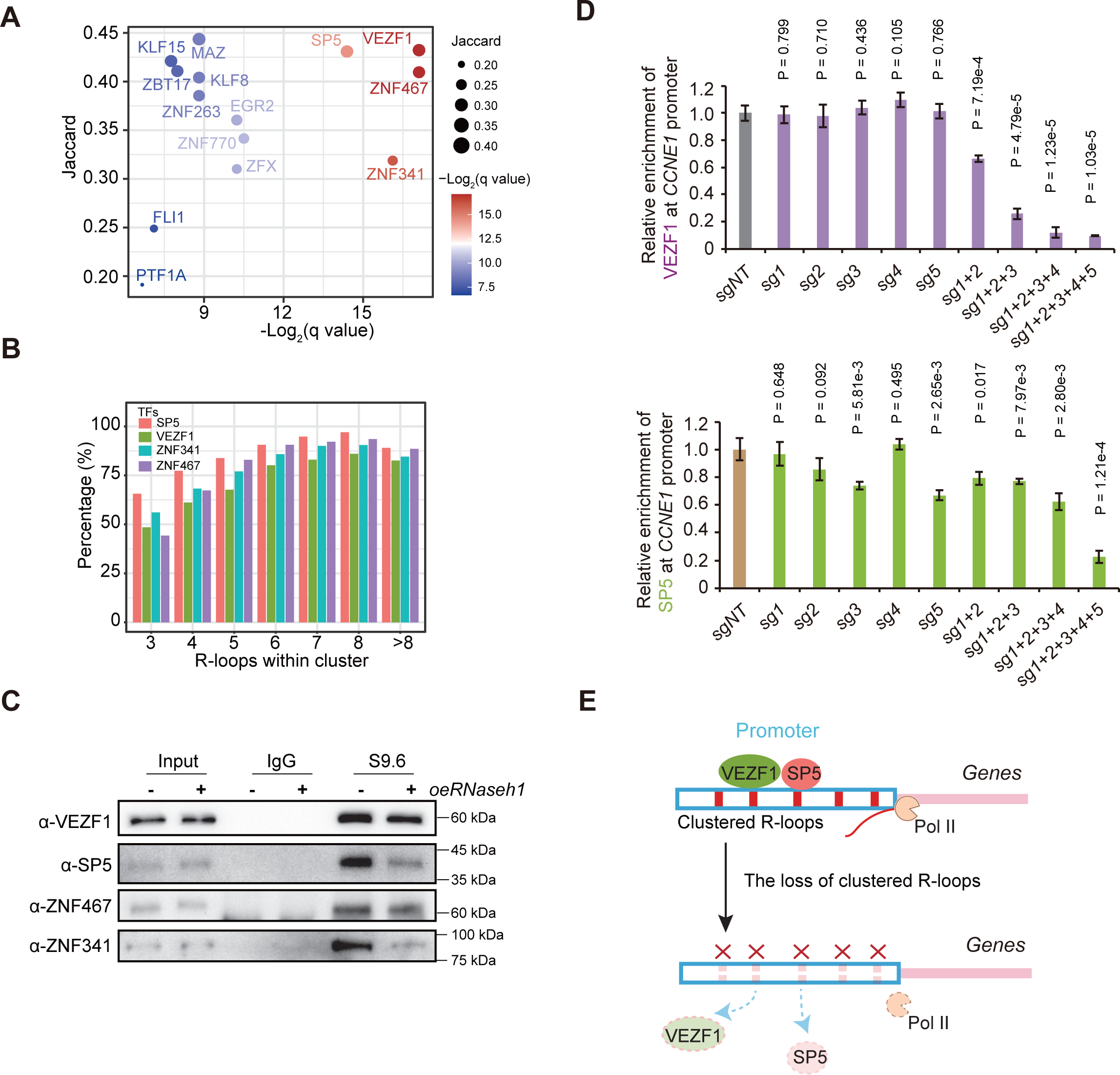
Loss of clustered R-loops causes defective gene expression by reducing the binding of zinc finger transcription factors at promoters. **A.** Transcription factor motifs identified from clustered R-loops, with the colors indicating the enrichment of the binding motifs of transcription factors and the size indicating the overlapping ratios. **B.** The percentage of clustered R-loops enriched with the motifs for selected transcription factors at promoters. **C.** Immunoprecipitation experiments showing the interactions between transcription factors and S9.6 antibody. **D.** qPCR showing the enrichment of VEZF1 and SP5 at promoters of *CCNE1* after clustered R-loop digested by dCas9-RNase H1. **E.** The model demonstrating the proposed mechanism by which clustered R-loops recruit zinc finger transcription factors at promoters for active transcription.

To investigate whether R-loops form complexes with these factors (VEZF1, SP5, ZNF467, and ZNF341), we performed S9.6 immunoprecipitation experiments. We observed that immunoprecipitated VEZF1, SP5, ZNF467, and ZNF341 by the S9.6 antibody were dramatically decreased after *RNaseh1* overexpression (Figure 5C). We found that clustered R-loops with more diverse binding motifs of these transcription factors showed a higher degree reduction of R-loop signals upon *RNaseh1* overexpression (Figure S13C). The number of downregulated genes gradually increased with the increase of types of binding motifs for these transcription factors in clustered R-loops (Figure S13D). Therefore, we speculate that clustered R-loops at promoters might recruit these zinc finger transcription factors.

VEZF1 and SP5 are required for RNA polymerase II recruitment and transcriptional regulation (Gowher et al., 2012; Harrison et al., 2000; Tang et al., 2017). Consistent with the changes in gene expression (Figure 4C and S11A), the loss of clusters with 5 R-loops but not 3 or 4 R-loops led to a noticeable reduction in the enrichment of RNA polymerase II (Figure S14A). We then analyzed the effect of clustered R-loops on the enrichment of VEZF1 and SP5 at promoters. CUT&RUN qPCR indicated that targeting 5 R-loops within a cluster by dCas9-RNase H1 at *CCNE1* promoter led to more robust downregulation of enrichment for both VEZF1 and SP5, while dCas9-RNase H1 had relatively less effect on the enrichment of either VEZF1 or SP5 at the promoter of *JUP* or *PHACTR4* (Figure 4B, 5D and S14B). In addition, only all 5 R-loops within a cluster at *CCNE1* promoter resolved by dCas9-RNase H1 caused the most significant decrease of the binding of both VEZF1 and SP5 compared with less than 5 R-loops resolved within *CCNE1* promoter (Figure 4B and 5D). Interestingly, our data indicated that only a reduction in the binding of both SP5 and VEZF1 by more than twofold resulted in a remarkable decrease in *CCNE1* gene expression (Figure 4C and 5D). We conclude that clustered R-loops, particularly those with five or more R-loops, recruit zinc finger transcription factors to gene promoters and activate gene expression.

## Discussion

R-loops are involved in diverse biological processes and diseases, yet their biogenesis, genomic distribution, and functions remain incompletely understood. Precise mapping of cellular R-loops could improve our understanding of their conflicting roles in different contexts of biology. Previous R-loop mapping methods have advantages and disadvantages. However, the lack of consistency among previous methods has hindered progress in R-loop biology. Therefore, the consistency and accuracy of R-loop identification are urgent issues in this field.

By developing RIAN-seq, we address the above challenges by providing an unbiased approach to identify R-loops at base-pair resolution, revealing precise positions and sizes of R-loops. We identify orders of magnitude more R-loops than other studies, offering a more comprehensive view of genome-wide R-loops and avoiding inconsistency of R-loops among previous methods in R-loop biology.

Interestingly, clustered R-loops facilitate the binding of zinc finger transcription factors, such as VEZF1 and SP5, to gene promoters, thereby activating transcription (Figure 5E). However, the exact mechanisms by which R-loops recruit these factors are still unknown and require further investigation. Previous studies indicated many factors are involved in R-loop interactome(Cristini et al., 2018; Wang et al., 2018). We supposed that these transcription factors bound to different R-loops within clustered R-loops may synergistically regulate gene expression. Furthermore, transcription factors function redundantly for gene activation(Ji et al., 2023). Thus, the loss of a fraction of R-loops within a cluster may be rescued by the other remaining R-loops within the same cluster, resulting in little effect on the binding of zinc finger transcription factors.

Zinc finger transcription factors activate transcription depending on the coregulators with which they interact(Kaczynski et al., 2003). Therefore, we propose that clustered R-loops might orchestrate the binding of coregulators of zinc finger transcription factors for gene activation. Moreover, R-loops at promoters are associated with active chromatin signatures containing H3K4me3 and H3K27ac(Sanz et al., 2016), clustered R-loops may influence chromatin states to further fine-tune transcription factors binding.

In summary, this study offers an unbiased and highly sensitive method for genome-wide R-loop identification at unprecedented resolution, which effectively enables us to decipher distinct features and functions of R-loops in various biological processes. The biogenesis of clustered R-loops that we identified is still a mystery. Unveiling the turnover of clustered R-loops will be crucial for understanding the functional roles of clustered R-loops and their regulatory mechanisms on transcription.

## Methods and Materials

### Cell culture

HEK293T was maintained in DMEM medium supplemented with 10% FBS. mESCs were cultured in DMEM medium supplemented with 15% FBS, 1 mM sodium pyruvate, 1 mM nonessential amino acids, 1 × GlutaMAX, 0.1 mM 2-mercaptoethanol, 1000 U/mL leukemia inhibitory factor, 3 mM CHIR99021 and 1 mM PD0325901.

### RIAN-seq

The genomic DNA was purified from HEK293T or mES cells via phenol/chloroform extraction and ethanol precipitation, or genomic DNA extraction kit. 1 µg DNA was incubated with NEB restriction enzyme mix (Alu I, Mbo I, Mse I, Dde I), Nuclease P1, T5 exonuclease, Lambda exonuclease at 37°C for 2 hr. As controls, RNase H (NEB, M0297S) was added during the incubation stage. 0.2 µL of Proteinase (Thermo Scientific, EO0491) was added before incubation at 55°C for 30 min and 70°C for 30 min. 0.3 µL 1M Tris-HCl (pH 8.0), 0.15 µL 1M MgCl_2_, 1.5 µL PEG200, 4.8 µL 50% PEG8000 and 1 µL transposase (Vazyme, TD501) were added and incubated at 55°C for 10 min. The transposition was stopped by the addition of 5 µL 0.25% SDS and incubation for 5 min at room temperature. Transposed DNA was gap-filled by addition of 2 µL Bst3.0 DNA polymerase (NEB, M0374S) and 25 µL NEBNext Q5 HotStart HiFi PCR Master Mix (NEB, M0543S). The mixture was incubated at 72°C for 15 min and 80°C for 10 min. For library amplification, the mixture was mixed with a barcoded i5 primer and a barcoded i7 primer PCR was performed to amplify the libraries using the following PCR conditions: 98°C for 45 s; thermocycling for 10 cycles at 98°C for 15 s, 65°C for 75 s; followed by 65°C for 5 min. The final libraries were purified with the 1 × AMPure beads (Beckman, A63882) size selection and were subjected to next-generation sequencing.

### Synthetic control experiment of RIAN-seq

Synthetic control ssDNA, ssRNA, dsDNA and RNA:DNA hybrid were mixed together at the indicated ratio used for RIAN-seq. After treatment, the level of synthetic oligos were assessed by qPCR.

### Real-time qPCR of RIAN-seq

The synthesized RNA:DNA hybrid, dsDNA, ssRNA and ssDNA were mixed with genomic DNA and treated with the RIAN-seq conditions.

### Site-specific editing of R-loops

dCas9-RNase H1 expressing plasmid and gRNA expressing plasmid were co-transfected into HEK293T cells, following by treatment with puromycin at a final concentration of 2 μg/ml for two additional days. Cells were harvested for qPCR analysis of gene expression, R-loop levels and the enrichment of transcription factors. The sequence of gRNAs was listed in Table S3.

### Co-IP

The nuclear protein extracts were incubated with 2 µg of S9.6 antibody or immunoglobulin G (IgG) and 25 µl of Dynabeads Protein A or Protein G (Invitrogen) in IP buffer [20 mM tris-HCl (pH 7.4), 150 mM KCl, 0.1% Triton X-100, and 5 mM EDTA (pH 8.0)] at 4°C for 12 hours. Beads were washed five times with IP buffer and then were boiled in 1× SDS loading buffer for the Western blot. The antibodies used in this study are as follows: VEZF1 antibody (Santa Cruz, sc-365560), SP5 antibody (Abcam, ab209385), ZNF467 antibody (BBI, D161356), ZNF341 antibody (Biorbyt, orb2276184).

### Sequencing data processing of RIAN-seq

Raw reads were processed using FastQC (v0.11.9) for quality control, and then low-quality base and adapter sequences were trimmed by Trim_galore (v0.6.7). The reads generated by this method were aligned to human (hg38) or mouse (mm10) reference genome using Bowtie2(Langmead and Salzberg, 2012) (v2.2.5). Reads of low mapping quality were filtered out utilizing SAMtools(Li et al., 2009) (1.14) with the parameters ‘-F 1804 -f 2 -q 20’. Concurrently, potential PCR duplicates were eliminated using Sambamba(Tarasov et al., 2015) (v0.7.1). The Bigwig files were generated by DeepTools(Fidel et al., 2016) (v3.5.1) with the parameter ‘-normalizeUsing RPKM -- extendReads 200’. In the generation of bulk cells, a binsize of 1 was employed, whereas a binsize of 10 was utilized for low-input conditions. The files were visualized in the Integrated Genomics Viewer (IGV) browser(James et al., 2011) and WashU Epigenome Browser(Zhou X Fau - Maricque et al., 2011). Picard tools were utilized to perform random sampling of reads, stratified by varying sequencing depths. Heatmaps and pile-up distributions across the genome, derived from various R-loop technologies or distinct treatments, were visualized using DeepTools(Fidel et al., 2016) (v3.5.1).

### Spike-in DNA normalization of RIAN-seq

The procedures for quality control and filtering of the raw reads were performed as above. The processed reads were then aligned to either the human (hg38)/mouse (mm10) or the Drosophila (dm6) (spike-in) reference genome using Bowtie2(Langmead and Salzberg, 2012) (v2.2.5). All unmapped reads and PCR duplicates were removed. For the results both with and without RNase H treatment, the number of reads on each base was quantified and subsequently normalized by the total number of spike-in reads (calculated as the total number of mapped spike-in reads divided by a constant).

### Peak calling for different R-loop mapping methods

DRIP-seq, DRIPc-seq, R-ChIP, R-loop CUT&Tag, MapR and spKAS-seq data in HEK293 cells were downloaded from GEO database. Other techniques were not evaluated mainly due to the lack of data corresponding HEK293T cells. All R loops were aligned to the human reference genome (hg38) using Bowtie2(Langmead and Salzberg, 2012) (v2.2.5). Given the diverse peak distributions exhibited by various R-loop techniques, we will employ distinct software and parameters tailored to each technique for peak calling. For the extremely wide techniques (DRIP-seq and DRIPc-seq), we used Epic2(Stovner and Saetrom, 2019) (v0.0.52) software and set --bin-size to 50 to call peaks. For the wide (spKAS-seq and MapR) and narrow (R-ChIP, R-loop CUT&Tag and this method) techniques, we used MACS2(Zhang et al., 2008) (v2.2.9.1) software and set different parameters to call peaks. Q-value ≤ 0.01 and fold change ≥ 5 were applied to broad and narrow peaks. Peaks with RPKM values greater than 30 were considered as high-confidence peaks and finally merged between multiple replicates. Peaks were annotated using clusterProfiler (4.10.1). For low input cells, we employed the MACS2 software to identify peaks with parameters set to -g hs --nomodel --nolambda. For an input of 1,000 cells, a Q-value ≤ 1e-3 was utilized. For inputs of 100 and 10 cells, a Q-value ≤ 1e-5 was applied. Peaks with RPKM values exceeding 100 were deemed to be of high confidence.

### Correlation analysis

Correlation calculations between replicate data were performed using the deepTools software package. First, multiBigwigSummary was used to calculate averaged read coverage within equally sized 10 kb bins of the entire genome. Regions in the human genome blacklist were excluded from the read coverage calculation. PlotCorrelation was subsequently used to calculate pairwise Pearson correlation coefficients with the output of multiBigwigSummary. Correlation calculations between low-input cells and bulk cells were performed using a similar approach but were based on gene-coding regions.

### GC skew and AT skew analysis

GC/AT skew was calculated using a 200 bp sliding window and a 10 bp step size as follows: GC skew = (G - C) / (G + C), AT skew = (A - T) / (A + T).

### RNA-seq analysis

Total RNA was extracted with TRIzol. An RNA library was generated with Vazyme mRNA-seq V3 Library Prep Kit for Illumina (Vazyme, NR611). To analyze the expression of genes, RNA-seq raw reads were trimmed using Trim_galore (v0.6.7) and aligned to the mouse genome (mm10 and Gencode gene annotation vM25) using STAR(Dobin et al., 2013) (v2.7.0d) and RSEM(Li and Dewey, 2011) (v1.3.3); The edgR(Robinson et al., 2010) package was used for differential expression analysis.

### Nascent RNA analysis

For the analysis of nascent RNA, we employed Trim_galore (v0.6.7) for adapter removal and pruning of low-quality reads. The reads were then aligned to the hg38 genome using Bowtie2 (v2.2.5), with alignments to ribosomal reads and PCR repeats being excluded. Peak calling was performed using MACS2 (v2.2.9.1).

### Signal to noise analysis

To compute the signal-to-noise ratio, we first determined the signal values for the enriched regions across various techniques. Subsequently, we employed BEDTools(Quinlan and Hall, 2010) (v2.31.0) to randomly sample bins of equivalent size and quantity to the peaks, serving as the genomic background. The final signal-to-noise ratio was then derived from the ratio of the signal values in the enriched regions to their corresponding random regions.

The FRiP score was derived by dividing the total count of reads situated within the peak region by the overall number of reads aligned to the reference genome. Fingerprints corresponding to various methods were visualized using DeepTools (v3.5.1).

For the construction of the ROC curve at genomic promoter sites, we annotated peaks within a 3000 bp range upstream and downstream of the genomic TSSs, designating these as positive sites, while the inverse was defined as negative sites. Owing to the variance in signal distribution across different R-loop technologies, we segmented the gene promoter locus into bins of 200 bp. The bin with the highest signal value within each promoter was computed as signal value from promoter regions. Ultimately, the ROC curve was plotted using plotROC(Sachs, 2017) (v 2.3.2).

### Identification of R-loop clusters

To find clustered R-loops, we used 1500 bp as the sliding window size and 10 bp as the sliding step to search R-loops from whole genome, the windows containing more than three R-loops were selected out. The overlapped windows were merged together to form the final R-loop clusters.

### Motif analysis

The MEME program(Bailey et al., 2009) was used to identify the enriched motifs. Using the generated count matrices, we used FIMO (MEME suite) to search for R-loops, the predicted motifs from MEME were matched to known motifs using TOMTOM.

## Data availability

All data have been deposited in the NCBI Gene Expression Omnibus (GEO) and are publicly available as of the date of publication. The previously published data are available under accession numbers GSE70189 (DRIP-seq, DRIPc-seq), GSE156400 (R-loop CUT&Tag), GSE192822 (spKAS-seq), GSE120637 (MapR), GSE97072 (R-ChIP), GSE112379 (PRO-seq), GSE139420 (KAS-seq).

## Acknowledgments

We thank the support from the biocomputing facility at Guangzhou National Laboratory. This work was supported by the National Key R&D Program of China (2021YFA1100300), and the National Natural Science Foundation of China (31925009, U21A20195, 32200456).

## Author contributions

H.Y. and Y.L. conceived and designed this project. Y.L. developed and performed the most experiments with the help of C.D. Y.S. conducted bioinformatics analysis. H.Y. and Y.L. wrote the manuscript with the input from Y.S. H.Y. supervised the entire study.

## Supplementary Figure legends

**Figure S1.**
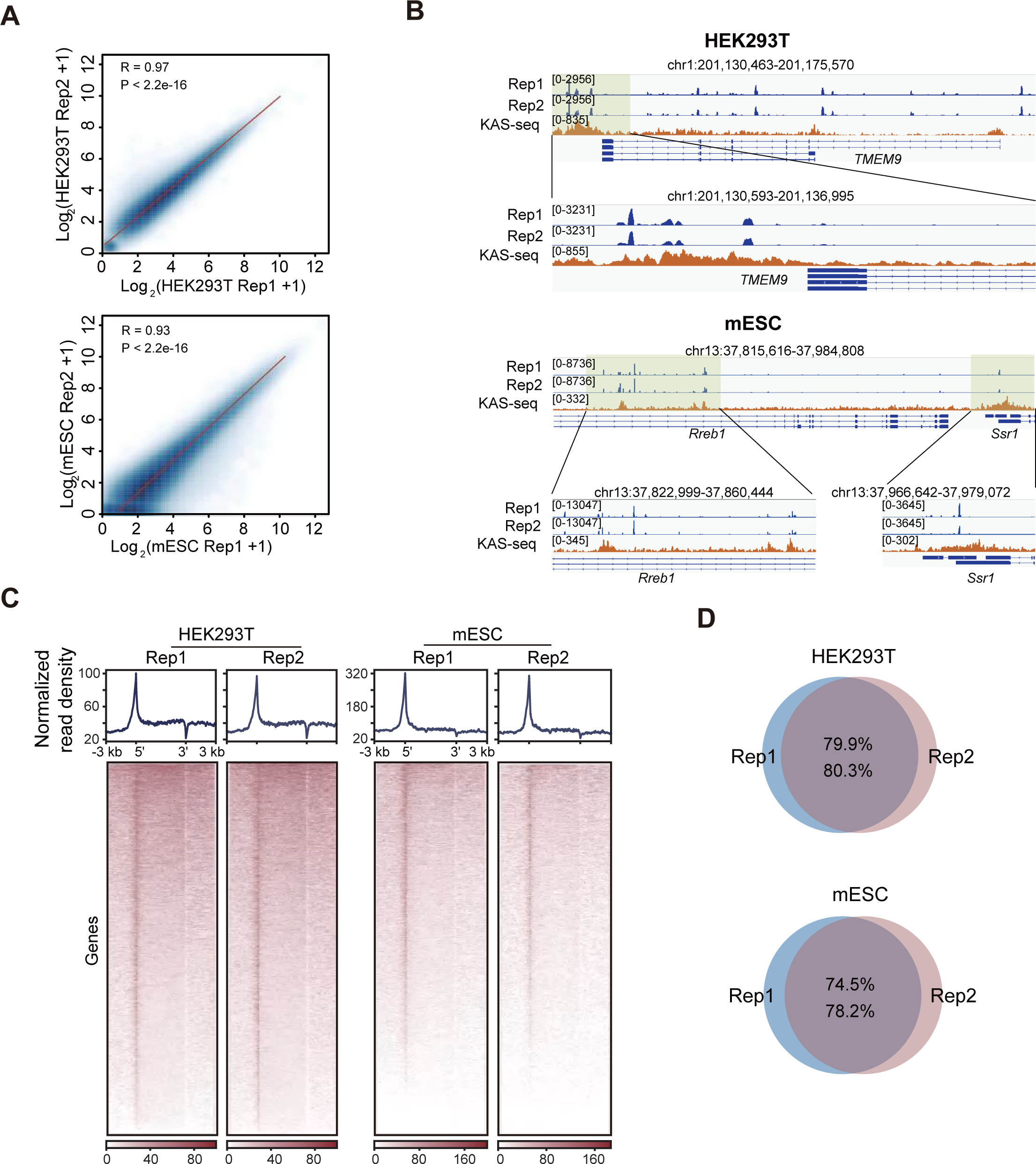
RIAN-seq generates reproducible data from biological replicates. **A.** The Pearson’s correlation between two replicates of RIAN-seq in HEK293T and mES cells. **B.** Genome browser snapshots depicting the signal of RIAN-seq and KAS-seq in HEK293T and mES cells. **C.** The metagene profiles of R-loops of two biological replicates across gene-coding regions in HEK293T and mES cells revealed by RIAN-seq. **D.** The overlapped R-loops between two replicates detected by RIAN-seq using HEK293T and mES cells.

**Figure S2.**
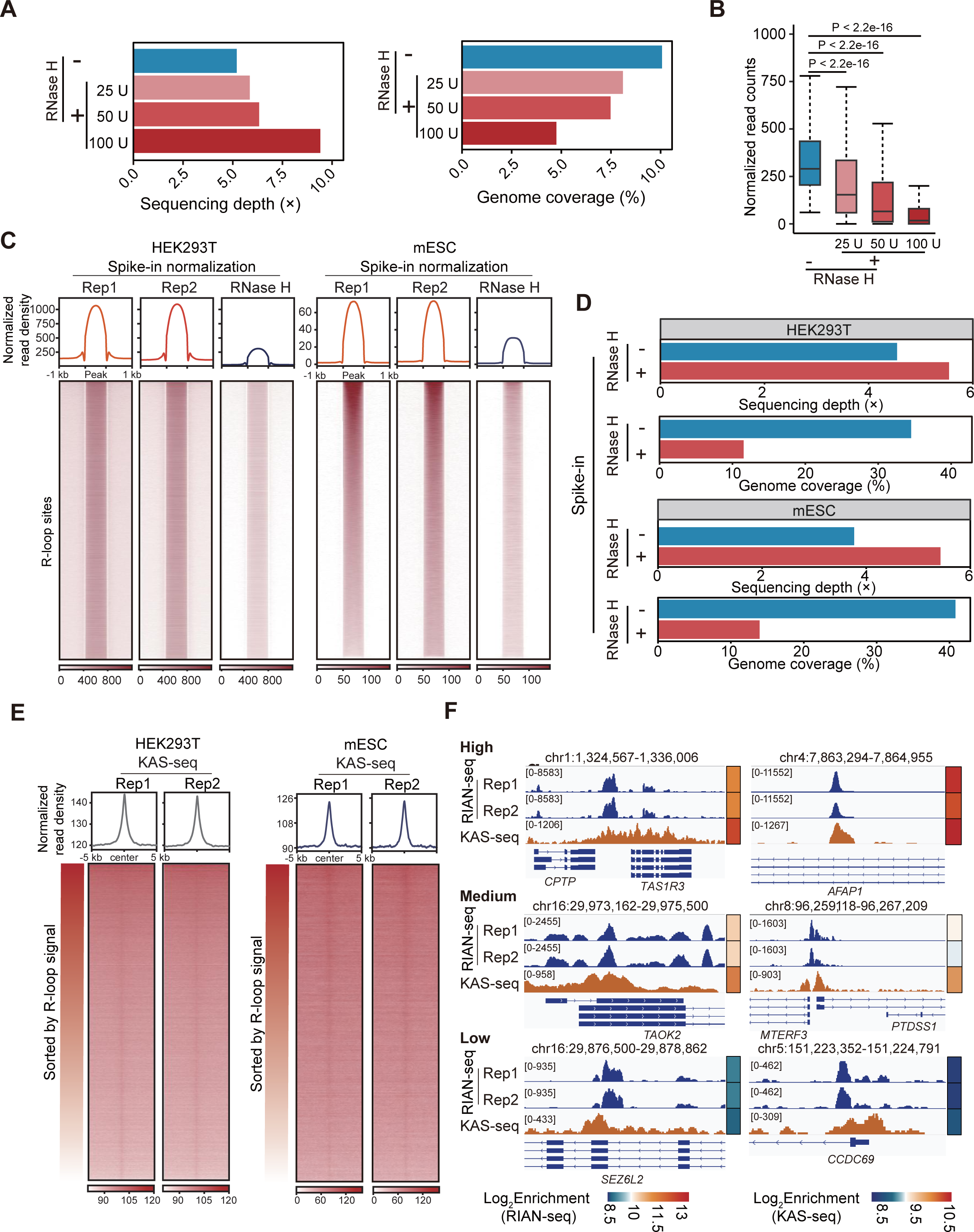
The signals of RIAN-seq can be resolved by RNase H. **A.** The bar graphs showing the sequencing depth and genomic coverage of R-loop signals in HEK293T cells treated with or without different dosages of RNase H using downsampled reads. **B.** The box plots depicting the density of R-loops identified by RIAN-seq treated with or without RNase H. **C.** The metaplots and heatmaps illustrating the signal density of RIAN-seq in HEK293T and mES cells with or without RNase H treatment. **D.** The bar graphs showing the sequencing depth and genomic coverage of R-loop signals in HEK293T cells treated with or without 25 U RNase H, normalized by spike-in. **E.** The metaplots and heatmaps showing the KAS-seq signal density of R-loops identified by RIAN-seq in HEK293T and mES cells. **F.** Snapshots indicating selected R-loops with high, medium, and low RIAN-seq signals alongside KAS-seq signals in HEK293T cells.

**Figure S3.**
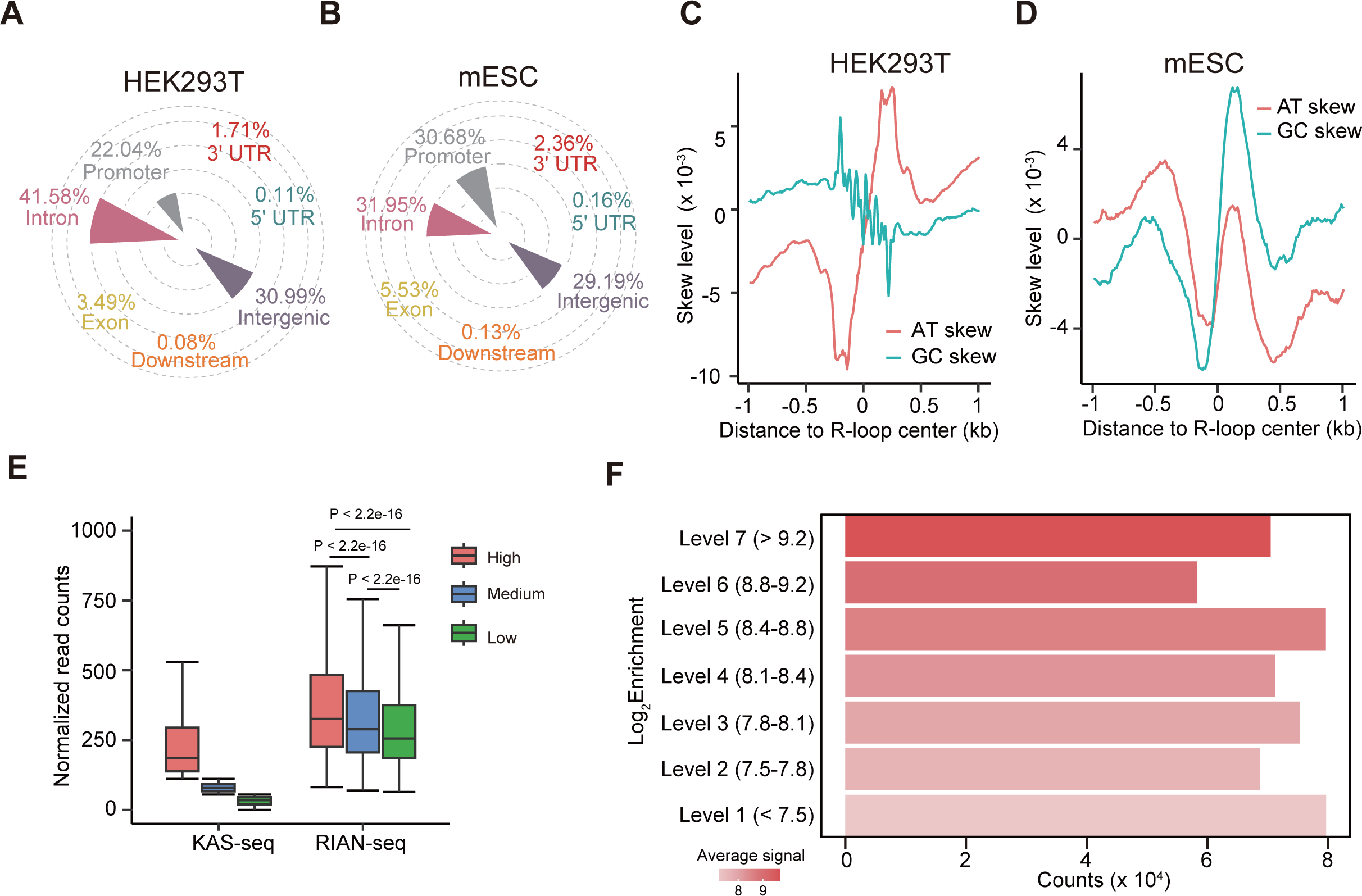
RIAN-seq effectively captures R-loops. **A.** The genomic distribution of RIAN-seq peaks in HEK293T cells. **B.** The genomic distribution of RIAN-seq peaks in mES cells. **C.** Metaplot showing AT and GC skew centered on peaks of RIAN-seq in HEK293T cells. **D.** The metaplot showing AT and GC skew centered on RIAN-seq peaks in mES cells. **E.** Box plots illustrating the density of high, medium, and low levels of R-loops identified by RIAN-seq, along with ssDNA signals captured by KAS-seq. **F.** The R-loop counts identified by RIAN-seq among different signal levels.

**Figure S4.**
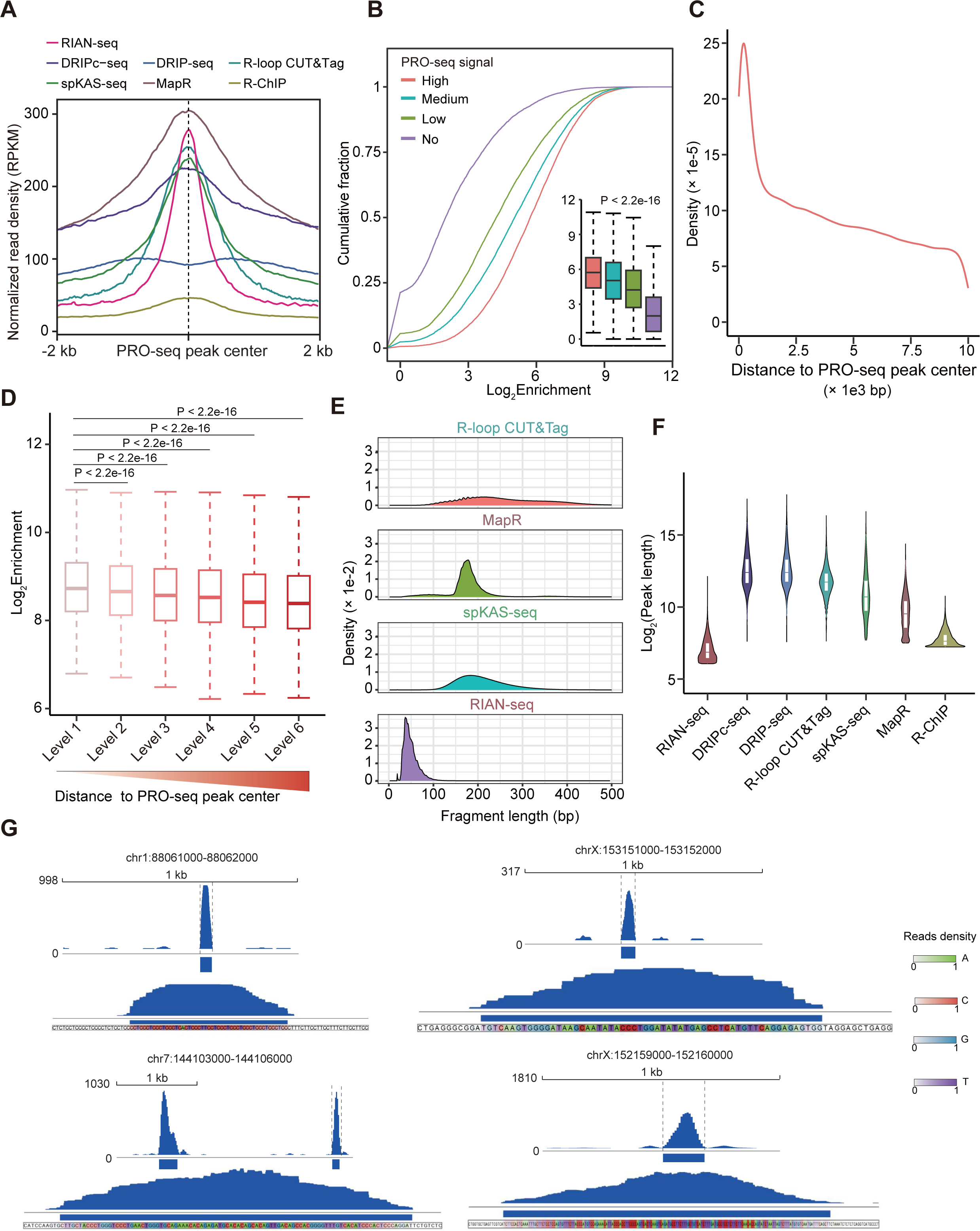
RIAN-seq attains base-pair resolution for R-loop detection. **A.** Density plots illustrating the distribution of peaks annotated using different R-loop mapping methods flanking PRO-seq peaks. **B.** Cumulative curves and boxplots illustrating the peak enrichment of RIAN-seq across four groups of PRO-seq peaks. **C.** Curve displaying the distribution of RIAN-seq peas flanking PRO-seq peaks. **D.** Box plots showing the signal level of RIAN-seq peaks situated within a certain absolute distance from PRO-seq peaks. **E.** The distribution of fragment length for RIAN-seq and other R-loop mapping method libraries. **F.** The length distribution of R-loops identified by RIAN-seq and other methods. **G.** Tracks indicating the width of signals of different R-loops identified by RIAN-seq.

**Figure S5.**
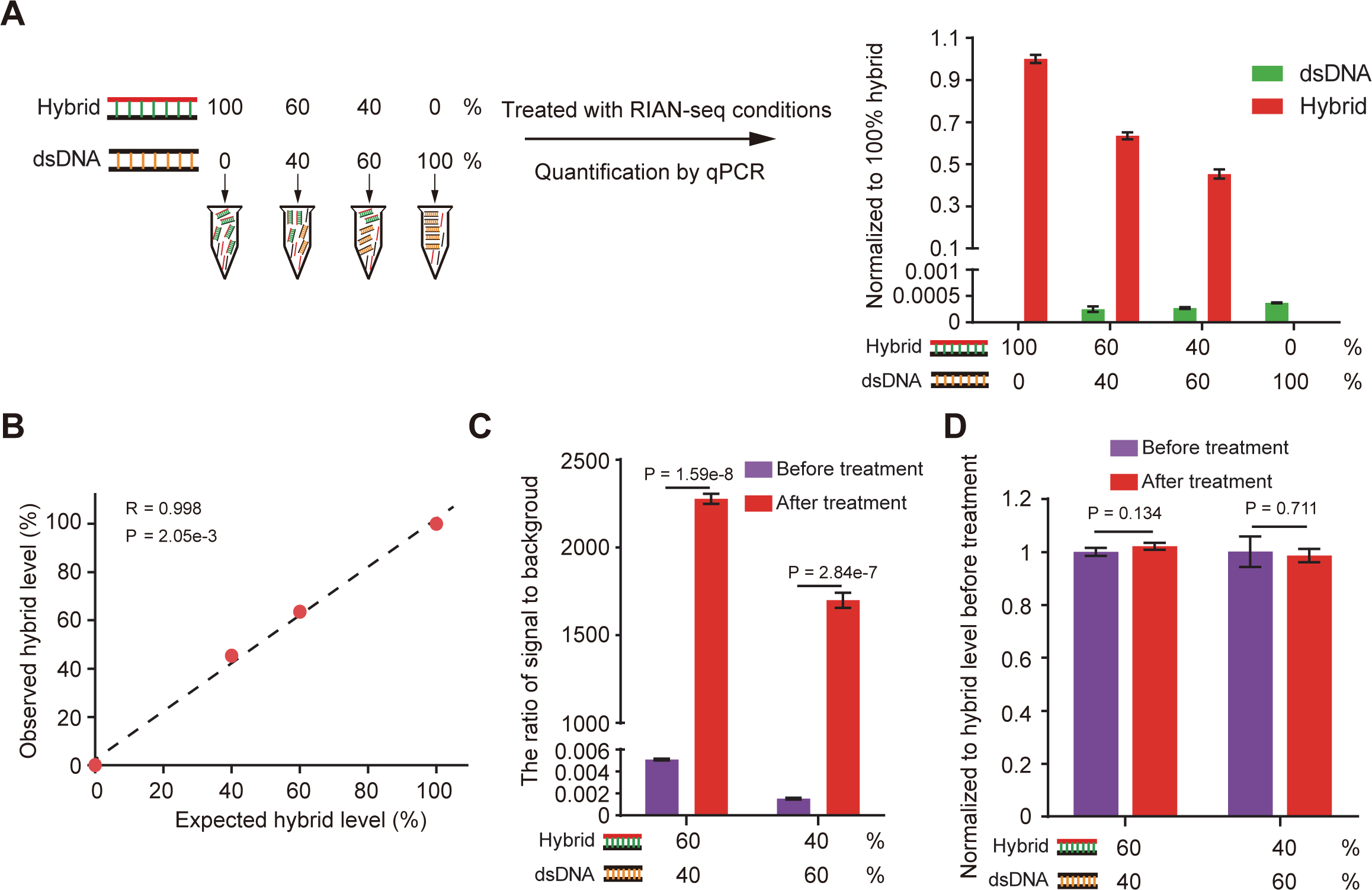
Assessment of specificity and background of RIAN-seq. **A.** Left, schematic illustration of the experimental setup for titration of synthetic oligos for RIAN-seq and qPCR assessment. Right, quantified levels of RNA:DNA hybrid and dsDNA following treatment of RIAN-seq. Data are presented as mean ± SD values. **B.** Correlation of expected and experimentally observed RNA:DNA hybrid levels based on the experiment outlined in **A**. The Pearson correlation coefficient and P-value based on a two-sided t-test are shown. **C.** qPCR assessment of RNA:DNA hybrid (signal) over dsDNA (noise) with two titrations of synthetic RNA:DNA hybrid and dsDNA. **D.** qPCR assessment of the RNA:DNA hybrid level before and after RIAN-seq experimental conditions with two titrations of synthetic RNA:DNA hybrid and dsDNA.

**Figure S6.**
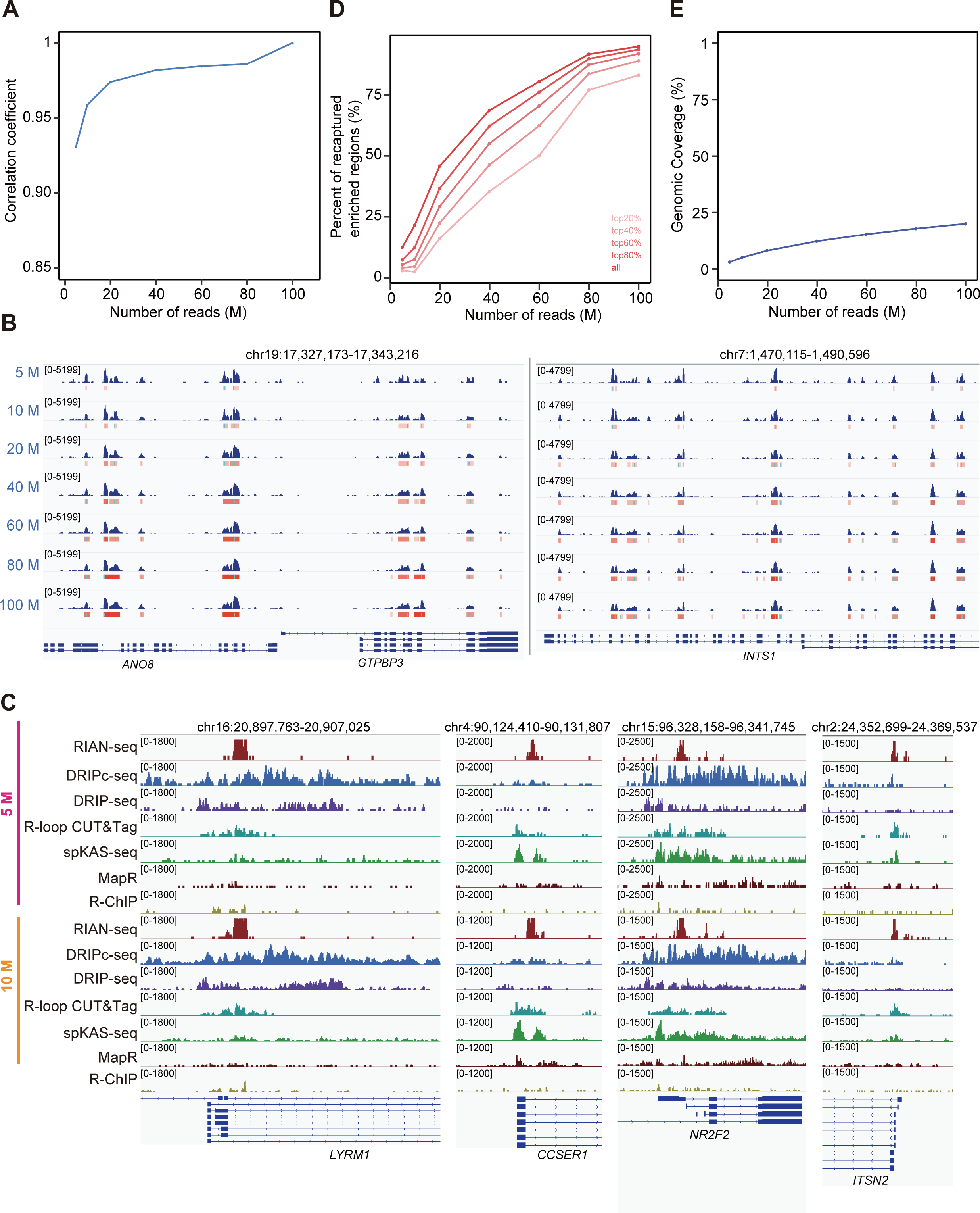
High-quality R-loop profiles of RIAN-seq. **A.** The Pearson’s correlation of signals across different sequencing depth data. **B.** The tracks depicting the signal distribution of RIAN-seq generated from different sequencing depths. **C.** The tracks generated from normalized sequencing depth showing the signal distribution and background from different R-loop detection methods. **D.** The percentage of captured R-loops via different sequencing depths of RIAN-seq. (E) Curve depicting the genomic coverage of R-loops identified by different sequencing depths of RIAN-seq.

**Figure S7.**
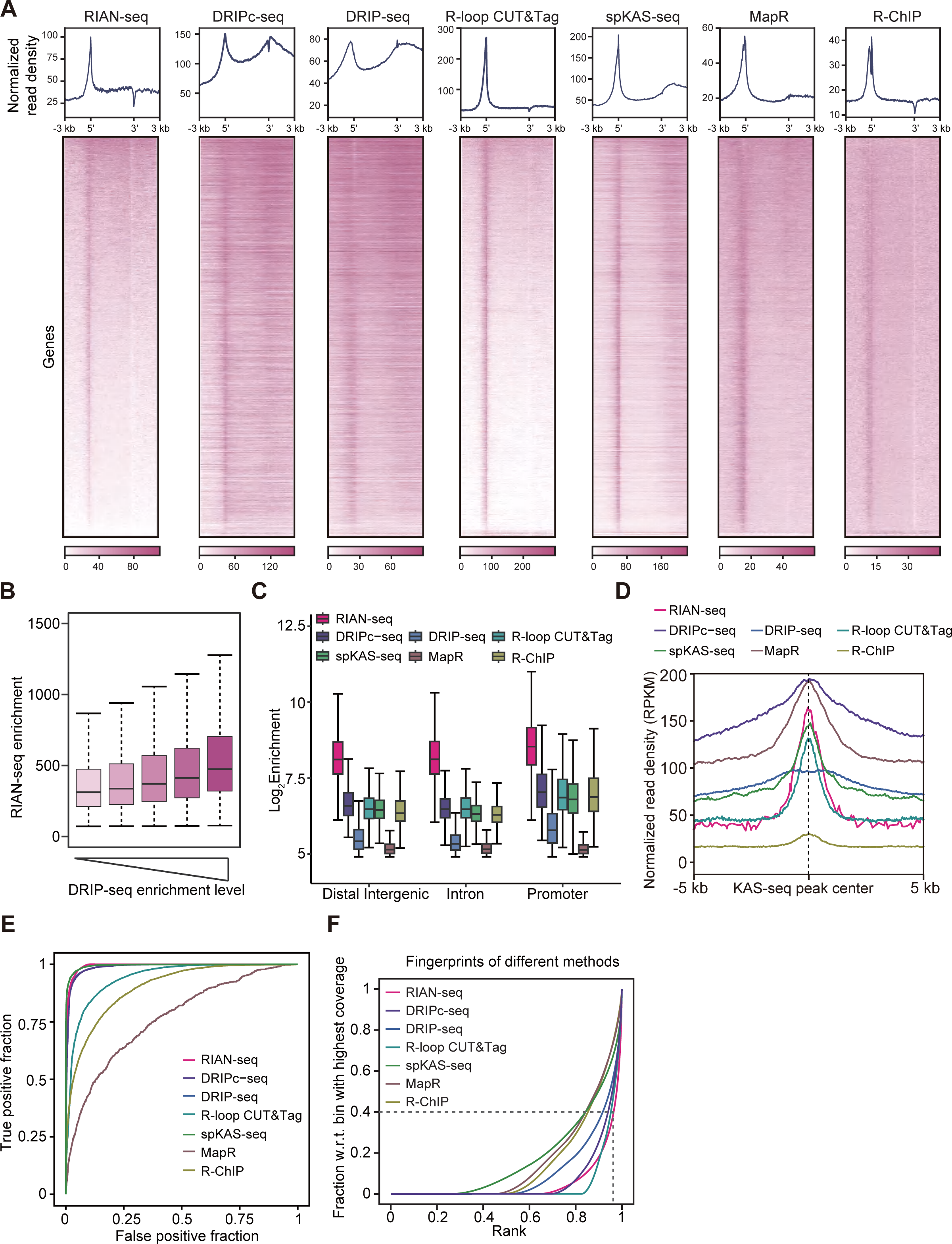
R-loop signal comparison between RIAN-seq and previously reported methods in HEK293T cells. **A.** The metagene plots and heatmaps depicting signals of different R-loop mapping methods at gene-coding regions. **B.** Box plots indicating that the signal enrichment of RIAN-seq is paralleled by DRIP-seq. **C.** The R-loop density at different genomic locations in different R-loop mapping methods. **D.** Density plots depicting the distribution of peaks of different R-loop mapping approaches flanking the peaks of KAS-seq. **E.** Receiver operating characteristics curves for sequencing data derived from different methods. **F.** Fingerprint plots of various R-loop mapping methods. w.r.t., with respect to.

**Figure S8.**
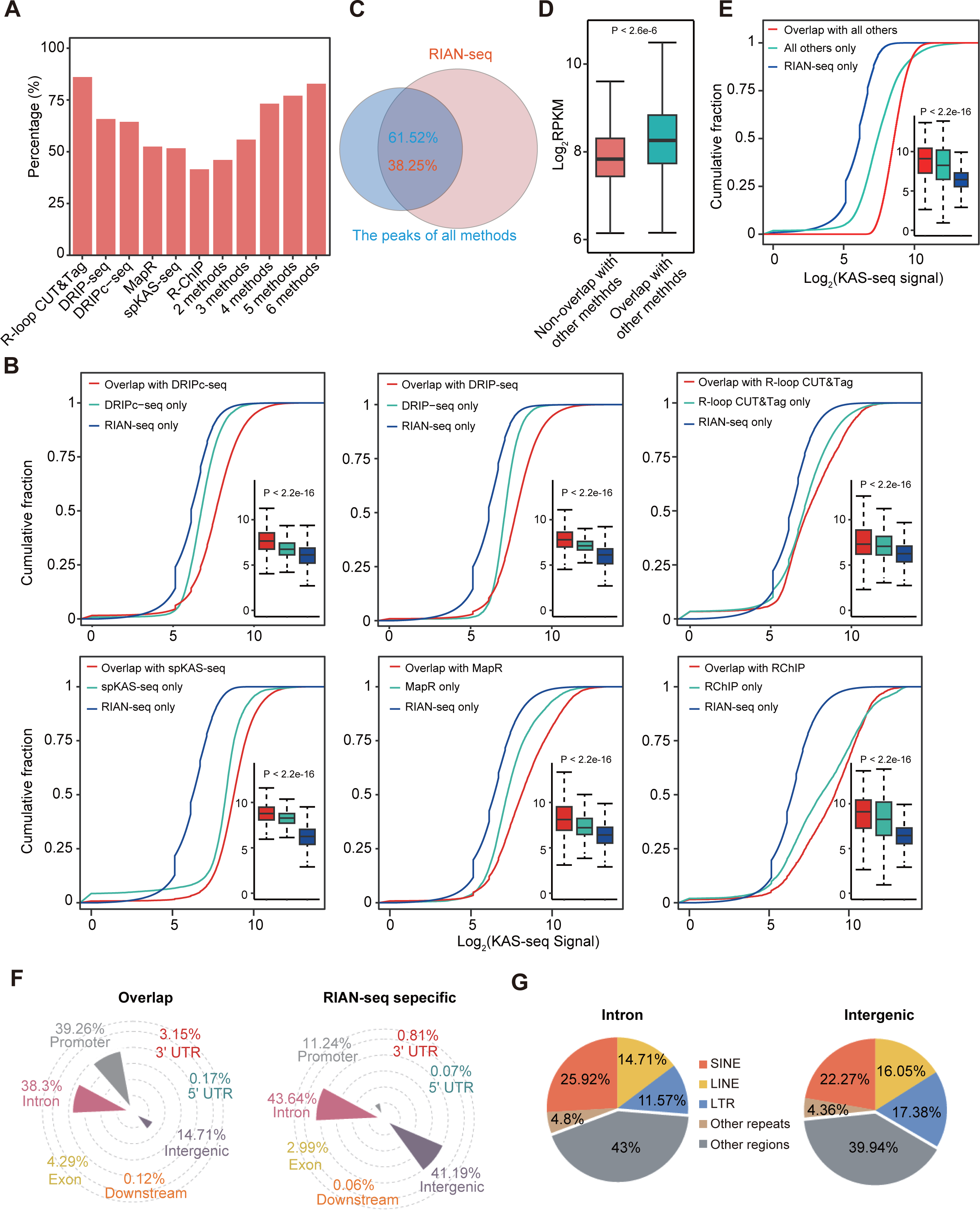
RIAN-seq captures most R-loops annotated by previous methods. **A.** The percentage of overlapped peaks between RIAN-seq and the other methods. The overlap rates between peaks were co-identified using two or more techniques, and RIAN-seq are also depicted. **B.** Cumulative distribution plots and box plots illustrating the ssDNA levels of overlapped R-loops and unique R-loops of different R-loop mapping methods. **C.** The percentage of overlapped peaks between RIAN-seq and all reported R-loops identified by the available methods. **D.** Box plot illustrating the signal density of overlapped R-loops and non-overlapped R-loops. **E.** Cumulative distribution plot and box plot depicting the ssDNA levels of overlapped R-loops and non-overlapped R-loops. **F.** The genomic distribution of overlapped R-loops and RIAN-seq specific R-loops. **G.** Pie charts depicting the detailed genomic feature distribution of intronic and intergenic R-loops unique in RIAN-seq.

**Figure S9.**
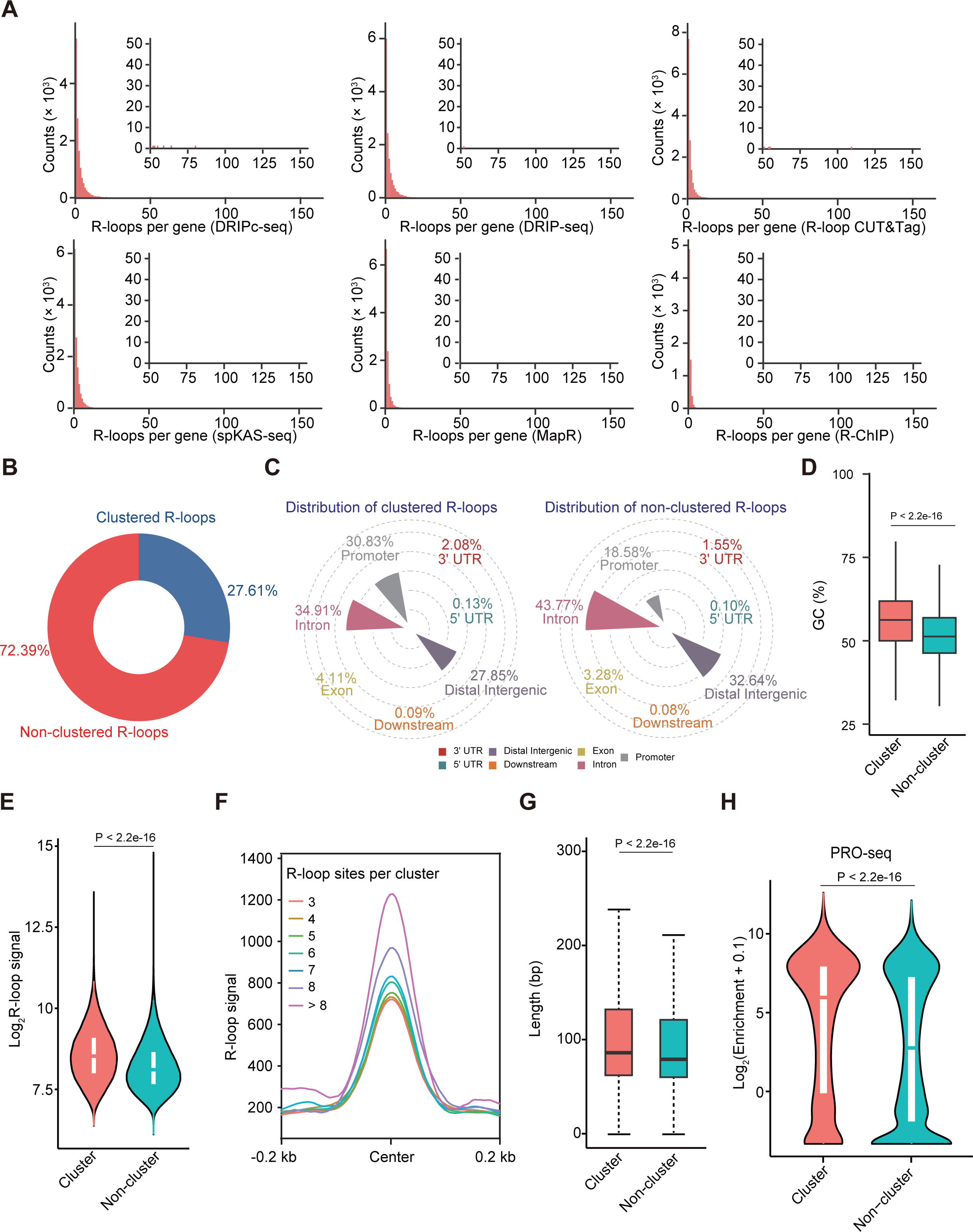
RIAN-seq identifies previously unknown distribution of R-loops. **A.** The number of genes containing R-loops identified by previous methods. **B.** The proportion of clustered R-loops. **C.** The genomic distribution of clustered R-loops and non-clustered R-loops. **D.** The comparison of GC content of clustered and non-clustered R-loops. **E.** The comparison of signal levels of clustered and non-clustered R-loops. **F.** R-loop signals of clusters containing different numbers of R-loops. **G.** Length distribution of clustered and non-clustered R-loops. **H.** Violin plot illustrating the expression level of genes with clustered and non-clustered R-loops at promoters using PRO-seq data.

**Figure S10.**
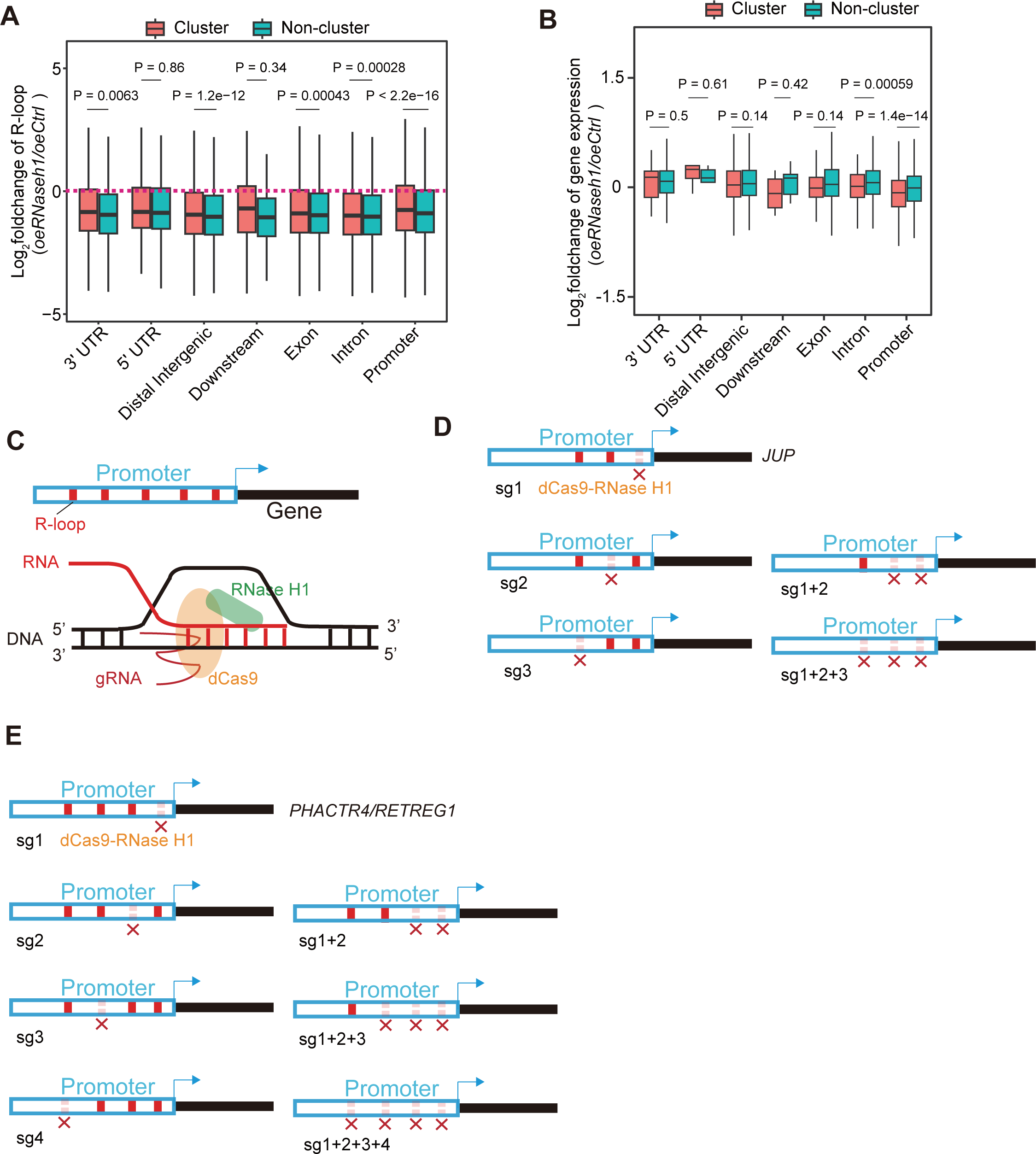
Overexpression of RNase H1 caused downregulated expression of genes with clustered R-loops. **A.** Box plots showing R-loop signal changes of genes with clustered and non-clustered R-loops, grouped by genomic distribution of R-loops. **B.** Box plots showing transcription changes of genes with clustered and non-clustered R-loops, grouped by genomic distribution of R-loops. **C.** The model showing site-specific R-loop edited by dCas9-RNase H1. **D.** The model showing the strategy of site-specific R-loop editing of 3 R-loops clustered in promoter of *JUP*. **E.** The model showing the strategy of site-specific R-loop editing of 4 R-loops clustered in promoter of *PHACTR4*, *RETREG1*.

**Figure S11.**
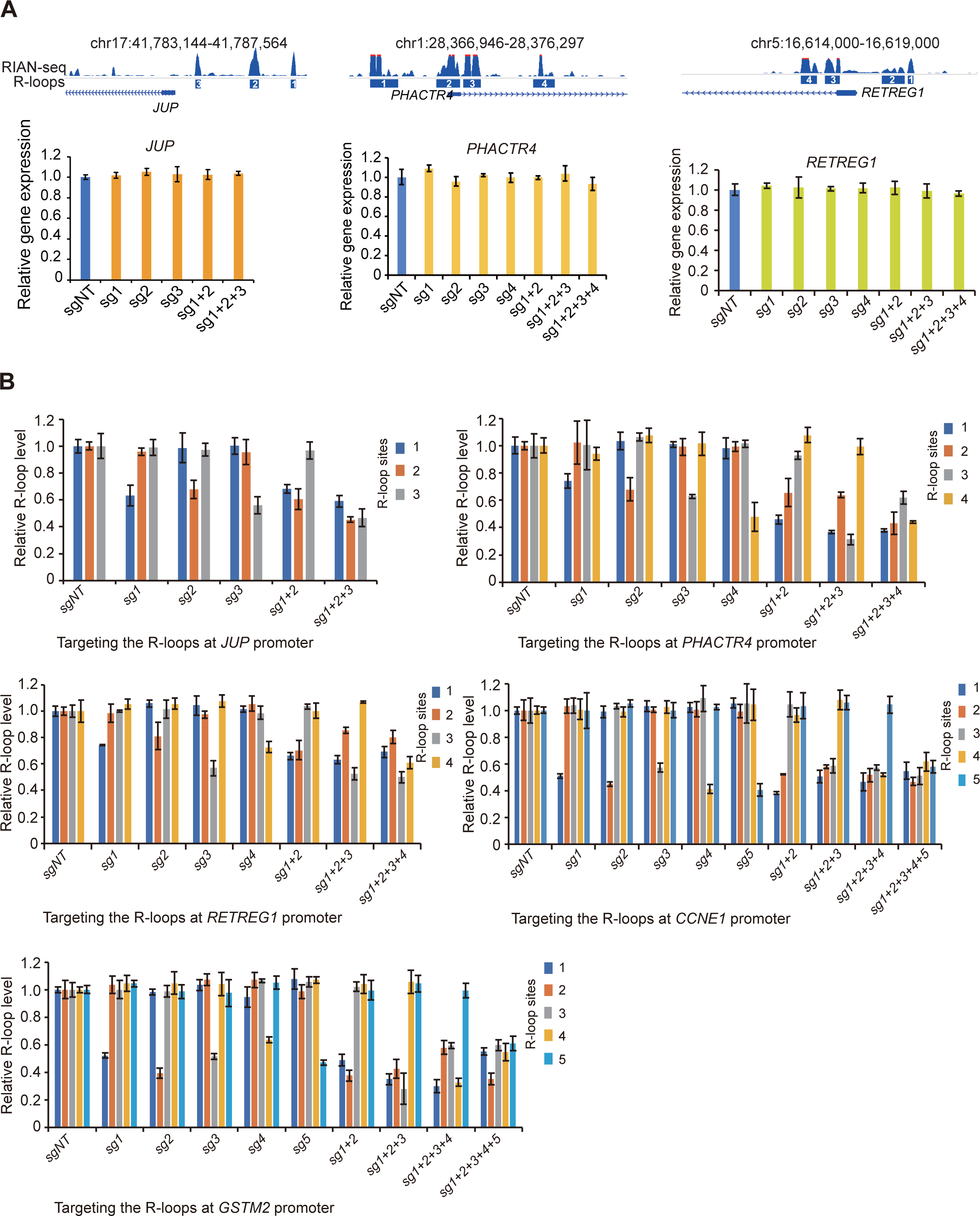
The changes of R-loops by dCas9-RNase H1 reduce gene expression. **A.** The bar graphs showing expression changes of genes with 3, 4 R-loops clustered in promoters after delivering dCas9-RNase H1 with different combination of gRNAs targeting R-loops. **B.** The bar graphs showing R-loop signal changes at promoters with 3, 4 and 5 clustered R-loops after delivering dCas9-RNase H1 with different combinations of gRNAs targeting R-loops.

**Figure S12.**
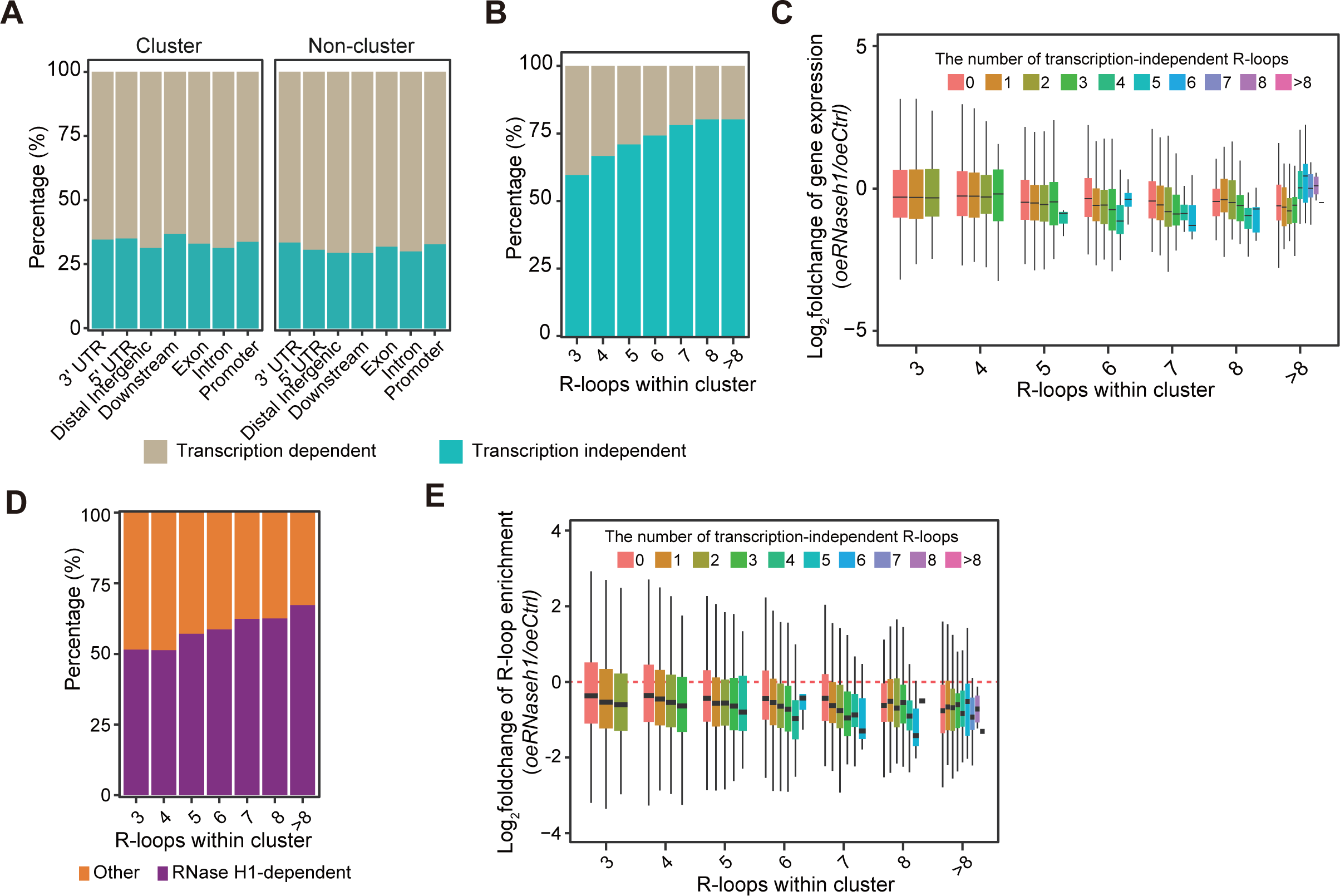
The interplay between clustered R-loops and transcription. **A.** The percentage of transcription-dependent and independent R-loops within clustered and non-clustered R-loops grouped by genomic distribution. **B.** The percentage of transcription-dependent and independent R-loop clusters grouped by different numbers of R-loops within a cluster in the promoter. **C.** The box plots showing expression changes of genes with different numbers of transcription-independent R-loops within clusters in the promoter. **D.** The percentage of transcription-independent R-loop clusters resolved by RNase H1 *in vivo*. **E.** The box plots showing R-loop signal changes at promoters with different numbers of transcription-independent R-loops within clusters.

**Figure S13.**
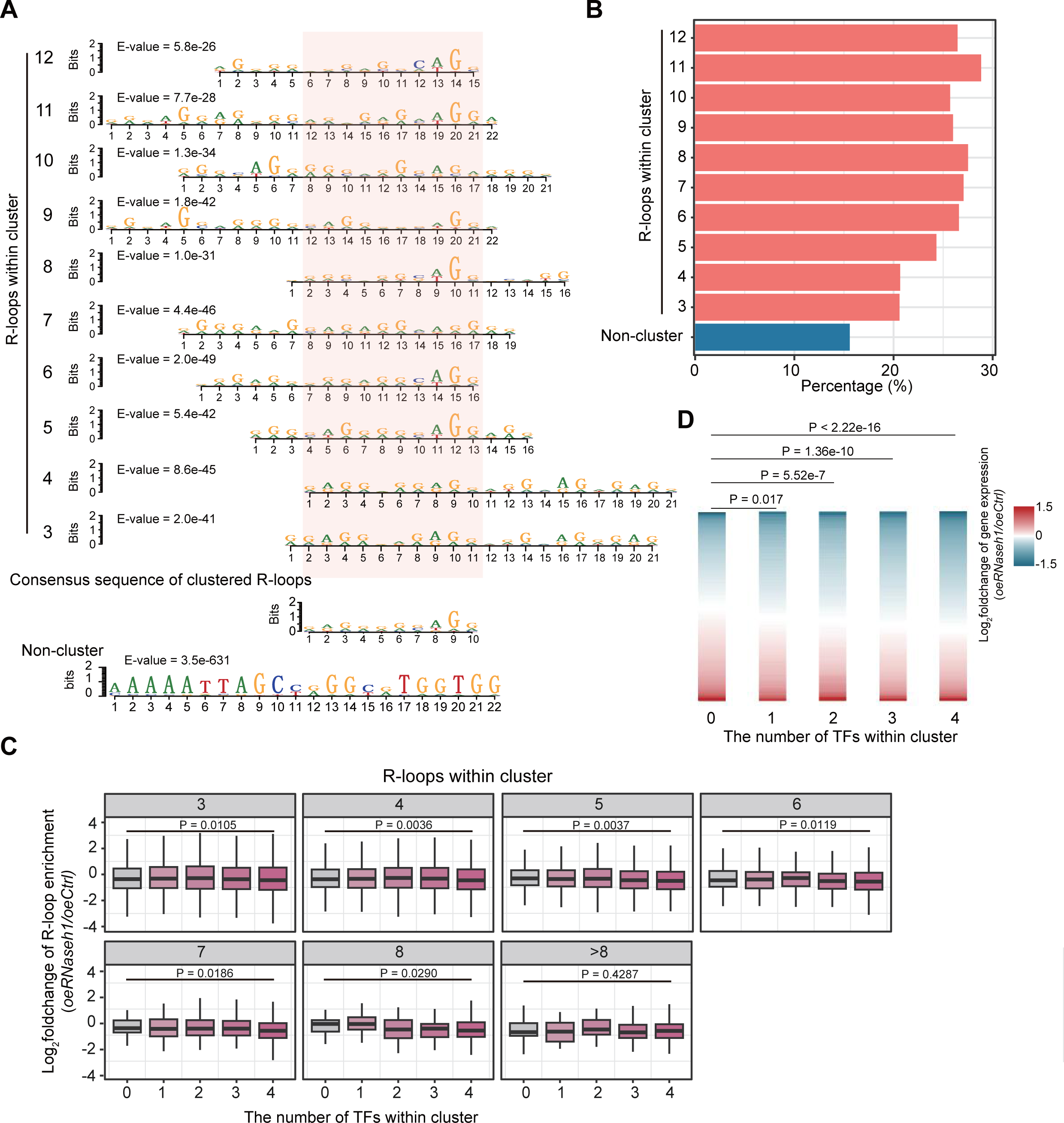
Clustered R-loops enrich binding motifs of zinc finger transcription factors. **A.** The enriched motifs among clustered and non-clustered R-loops. **B.** The percentage of consensus sequence of clustered R-loops in clustered and non-clustered R-loops. **C.** The box plots showing signal changes of R-loop clusters with different numbers of transcription-independent R-loops within clusters in promoter grouped by the number of types of transcription factors binding motifs. **D.** Heatmaps showing expression changes of genes with clustered R-loops in promoter grouped by the number of transcription factors binding motifs.

**Figure S14.**
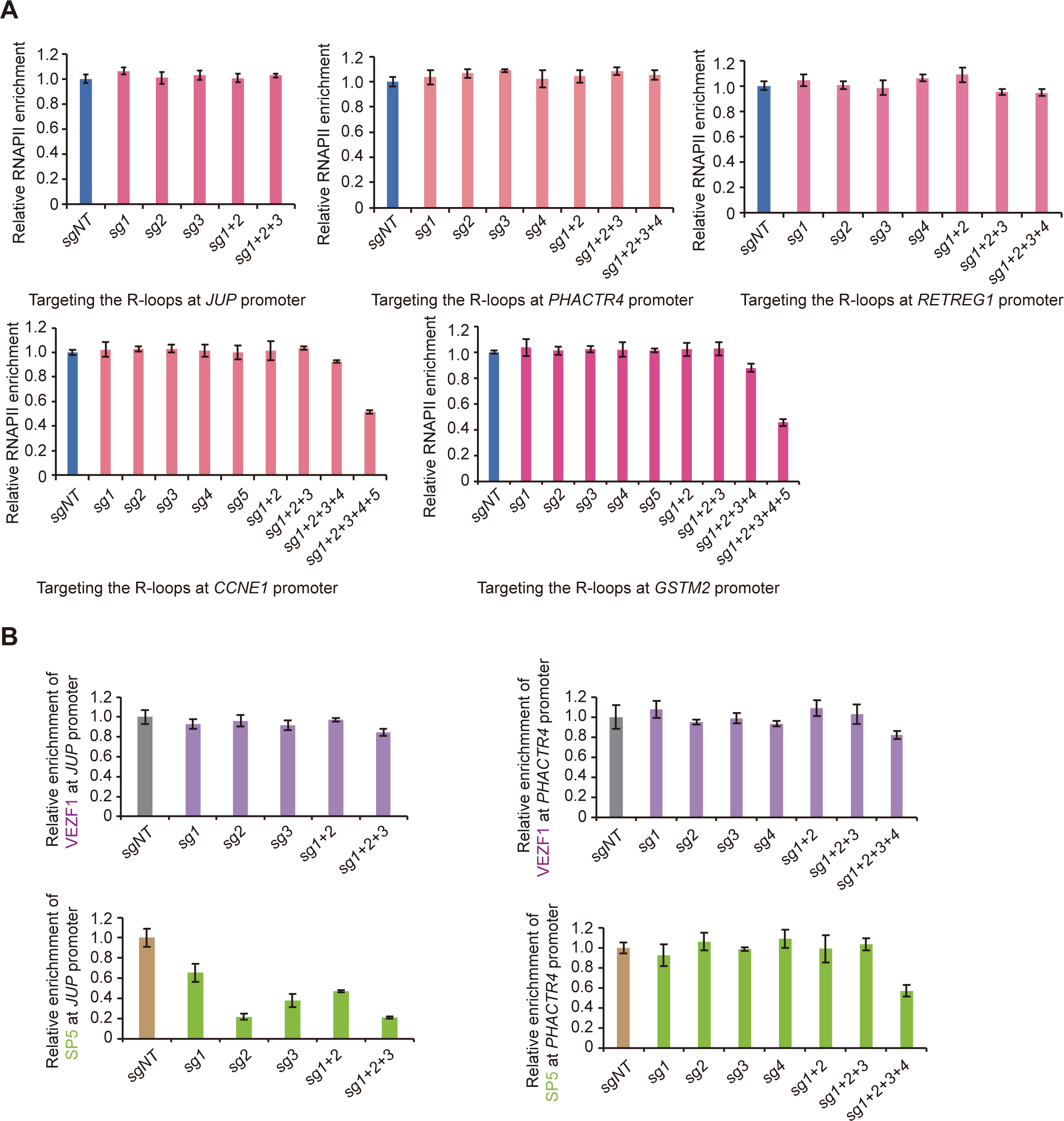
Clustered R-loops regulate the occupancy of RNA polymerase II and transcription factors at promoters. **A.** The bar graphs showing the enrichment of RNA polymerase II after treatment by dCas-9-RNase H1 and gRNAs targeting clustered R-loops in promoter of *JUP*, *PHACTR4*, *RETREG1*, *CCNE1*, *GSTM2*. **B.** CUT&RUN qPCR indicating the enrichment of VEZF1 and SP5 at promoters of *JUP* and *PHACTR4* after R-loops digested by dCas9-RNase H1.

